# Single-nuclei sequencing of skeletal muscle reveals subsynaptic-specific transcripts involved in neuromuscular junction maintenance

**DOI:** 10.1101/2024.05.15.594276

**Authors:** Alexander S. Ham, Shuo Lin, Alice Tse, Marco Thürkauf, Filippo Oliveri, Markus A. Rüegg

## Abstract

The neuromuscular junction (NMJ) is the synapse formed between motor neurons and skeletal muscle fibers. Its stability relies on the continued expression of genes in a subset of myonuclei, called NMJ myonuclei. Here, we use single-nuclei RNA-sequencing (snRNA-seq) to identify numerous undescribed NMJ-specific transcripts. To elucidate how the NMJ transcriptome is regulated, we also performed snRNA-seq on sciatic nerve transected, botulinum toxin injected and *Musk* knockout muscles. These data show that NMJ gene expression is not only driven by agrin-Lrp4/MuSK signaling, but is also affected by electrical activity and trophic factors other than agrin. By selecting three previously undescribed NMJ genes *Etv4*, *Lrtm1* and *Pdzrn4*, we further characterize novel contributors to NMJ stability and function. AAV-mediated overexpression and AAV-CRISPR/Cas9-mediated knockout show that *Etv4* is sufficient to upregulate expression of ∼50% of the NMJ genes in non-synaptic myonuclei, while muscle-specific knockout of *Pdzrn4* induces NMJ fragmentation. Further investigation of *Pdzrn4* revealed that it localizes to the Golgi apparatus and interacts with MuSK protein. Collectively, our data provide a rich resource of NMJ transcripts, highlight the importance of ETS transcription factors at the NMJ and suggest a novel pathway for NMJ post-translational modifications.

## Introduction

Skeletal muscle fibers form a syncytium, with many myonuclei sharing a single cytoplasm, resulting from the fusion of mono-nucleated progenitors. In mice, muscle fibers can reach several micrometers in length, accommodating hundreds of myonuclei ^1^. Although all myonuclei within a fiber share one cytoplasm, their gene expression repertoire is not homogenous. Based on location and gene expression signature, myonuclei can be categorized into three main groups ^2, 3, 4, 5^. The largest myonuclei category are distributed throughout the fiber and express a set of contractile and metabolic genes specific to their fiber type. We refer to this group as ‘body myonuclei’ ^2^ as they make up the “body” of the muscle fiber, but they have also been called bulk myonuclei ^4^ or simply named after the fiber type they belong to ^3^. Two substantially smaller but specialized populations, typically called myotendinous junction (MTJ) and neuromuscular junction (NMJ) myonuclei ^2, 3, 4, 5^, are localized at muscle-tendon and muscle-motor neuron contact sites, respectively. NMJ myonuclei, also known as subsynaptic or fundamental myonuclei ^6^, in mice consist on average of four nuclei clustered beneath the muscle postsynapse ^7, 8, 9^. They express transcripts whose products are enriched at the postsynapse and are often of functional importance for the NMJ. Examples are the acetylcholine receptor (AChR) subunits *Chrne*, *Chrna1*, *Chrnd*, *Chrnb1* ^10^, which encode the AChRε, AChRα, AChRδ and AChRβ subunits, respectively, and *Musk*, which encodes the muscle-specific kinase (MuSK) that is essential for NMJ formation and maintenance ^11, 12, 13^. This localized expression of NMJ genes allows postsynaptic enrichment of the requisite proteins to form and maintain the NMJ.

Spatial restriction of synaptic proteins to central regions of muscle begins during embryonic development, before motor axons reach the muscle fibers, a phenomenon known as prepatterning ^6, 14^. Innervation by motor neurons condenses and further enriches postsynaptic proteins, most likely *via* neural agrin release from motor axons. Neural agrin binds to the transmembrane protein LDL receptor-related protein 4 (Lrp4) expressed by muscle fibers ^15, 16^. As a result, Lrp4 clusters at the NMJ ^17^ where it forms a receptor complex with MuSK to promote intracellular cross-autophosphorylation and MuSK activation. Full activation of MuSK requires the adaptor protein downstream of tyrosine kinases-7 (Dok7), which additionally supports the recruitment of downstream proteins to the MuSK scaffold ^18^. Together, these proteins locally transduce the synaptic signal from neural agrin into muscle fibers to induce NMJ gene expression in nearby myonuclei that aggregate at the nerve-muscle contact site and become NMJ myonuclei ^19, 20^. In turn, release of acetylcholine (ACh) and the resulting action potentials in muscle fibers suppress expression of NMJ genes in body myonuclei ^10, 21^. Suppressed transcripts also include the developmental *Chrng* isoform while agrin-Lrp4/MuSK-signaling at the NMJ initiates the expression of the adult *Chrne* isoform ^6, 20^. Hence, there is a close interplay between local, “trophic” signals that maintain synaptic gene expression in NMJ myonuclei and “suppression” signals that spread throughout the muscle fibers to prevent synaptic gene expression in body myonuclei ^20^. The recent use of single-nuclei RNA-sequencing (snRNA-seq) has allowed to identify NMJ transcripts in an unbiased manner ^2, 3, 4, 5^. As there are still many neuromuscular disorders of unknown etiology, some of which show a “dying back” phenomenon, that can be ameliorated by overexpression of Dok7 ^22, 23, 24^, studying the transcripts expressed by NMJ myonuclei may also provide new insights into such diseases.

In this study, we used snRNA-seq to identify NMJ transcripts in adult mouse muscle. To examine how motor neurons contribute to the enrichment of NMJ transcripts in NMJ myonuclei we transected the sciatic nerve, which abrogates “trophic” support from the nerve and the “suppression” signals by electrical activity. To distinguish the effects from trophic support and electrical activity we injected botulinum toxin (BoTX), which prevents ACh release and hence specifically abrogates electrical activity. Additionally, to study which NMJ transcripts are enhanced through agrin-Lrp4/MuSK-signaling we muscle specifically knockout *Musk* and perform snRNA-seq before a prominent denervation signature occurs. From our snRNA-seq data we selected three NMJ transcripts and characterized them by overexpression and muscle fiber-specific knockout studies. We show that overexpression of the transcription factor ETV4 is sufficient to drive expression of NMJ transcripts in body myonuclei and that the Golgi-associated protein PDZRN4 affects AChR and MuSK localization at the postsynapse. Taken together, our data provide a rich resource of NMJ enriched transcripts, available for exploration on our online snRNA-seq atlas (https://ruegglab.shinyapps.io/snrnaseq/). Additionally, we provide new insights into the function of three previously undescribed proteins at the NMJ.

## Results

### Identification of transcripts expressed in NMJ myonuclei

To identify NMJ myonuclei-enriched transcripts, we developed a new single nuclei isolation protocol for skeletal muscle that bypasses the need for Fluorescence-Activated Cell Sorting (FACS). Single nuclei suspensions were prepared from two *tibialis anterior* (TA) and two *gastrocnemius* (GAS) muscles of adult C57BL/6 mice. Following barcoding, library preparation, sequencing and filtering, we generated a UMAP with ∼36’000 nuclei expressing an average of ∼1700 genes each (Fig. 1a). Using canonical cell type markers, we identified 11 mono-nucleated cell types and five myonuclei populations (Fig. 1b), similar to previous work ^2, 3, 5^. Our unbiased UMAP clustering generated two clusters that have not been described in previous skeletal muscle snRNA-seq datasets. By differential expression analysis, we identified *Mmrn1* and *Tenm2* as transcriptional markers for these two populations. Using single molecule fluorescence *in situ* hybridization (smFISH), *Mmrn1*-expressing nuclei localized near larger blood vessels, visualized by immunofluorescence (IF) staining against laminin-α4 ^25, 26^, while *Tenm2*-expressing nuclei were located close to motor neuron axons, visualized by IF staining against neurofilament (Supplementary Fig. S1a). As *Mmrn1*+ nuclei also express *Pecam1* (Fig. 1b), we concluded that they represent a subtype of endothelial cells. *Tenm2*+ nuclei were also identified in a single-cell sciatic nerve atlas ^27^, suggesting that they represent a peri- and endoneurial cell type.

**Fig. 1:**
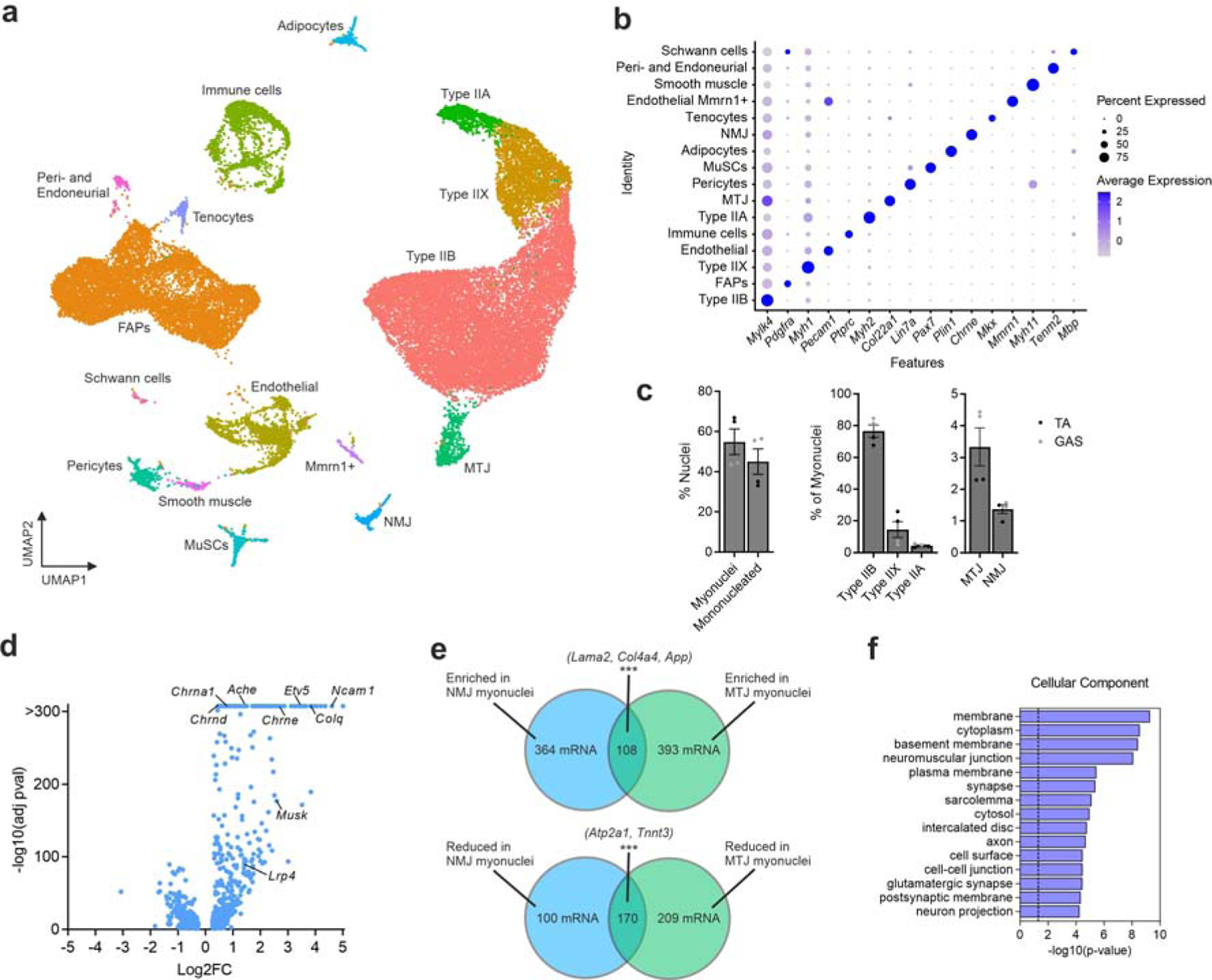
Identification of transcripts enriched in NMJ myonuclei. **a** UMAP of ∼36’000 nuclei isolated from *tibialis anterior* (TA; *n* = 2) and *gastrocnemius* (GAS; *n* = 2) muscles from four adult wild-type, male mice. **b** Dot-plot of marker genes used for cell/nuclear type identification. **c** Proportion of nuclei and myonuclei subtypes found in TA and GAS muscles. **d** Volcano plot visualizing differentially expressed transcripts in NMJ myonuclei compared to body myonuclei (max. padj. cutoff at 2.23E-308). Examples of known NMJ transcripts are indicated. **e** Venn diagram showing the number of transcripts enriched/reduced in NMJ/MTJ myonuclei and how they overlap (padj. < 5E-5, log2FC ±0.25). **f** Top 15 cellular component gene ontology (GO) terms associated with transcripts enriched in NMJ myonuclei (472 transcripts, padj. < 5E-5, log2FC > 0.25). GO-term analysis was performed with DAVID. In (**c**) data is presented as mean ± SEM. In (**f**) p-value = benjamini-hochberg p-value.

Myonuclei were identified based on the expression of *Ttn* and *Neb* and represented ∼45% and ∼65% of nuclei in GAS and TA muscle, respectively (Fig. 1c, Supplementary Fig. S1b). Within the myonuclei population type IIB, IIX and IIA myonuclei were distinguishable based on the expression of *Myh4*, *Myh1* and *Myh2*, respectively (Fig. 1c, Supplementary Fig. S1b). For type IIB myonuclei, we found other more striking gene markers such as *Mylk4* (Fig. 1b, Supplementary Fig. S1b). MTJ and NMJ populations gave rise to additional distinct clusters and made up ∼3% and ∼1% of myonuclei, respectively (Fig. 1c). The distinct clustering of these two small myonuclei populations highlights their unique transcriptional identity. Indeed, hundreds of genes were differentially expressed in the NMJ and MTJ populations compared to body myonuclei (Fig. 1d, Supplementary Fig. S1c). Overall, more genes were enriched, than depleted in these specialized populations, indicating they largely maintain their traditional myonuclear identity despite their additional specialized transcriptional responsibilities. In line with this notion, a higher transcriptional activity in NMJ myonuclei has previously been suggested based on chromatin organization ^28^. We also explored possible similarities between NMJ and MTJ myonuclei by comparing enriched and reduced transcripts from both populations (Fig. 1e). Many commonly enriched NMJ and MTJ myonuclei transcripts coded for membrane or extracellular matrix proteins (e.g. *Lama2*, *Col4a4* and *App*), indicating an increased need for locally produced structural proteins to support these crucial connection sites between the muscle fiber and motor axons and tendons, respectively. Conversely, several transcripts encoding proteins involved in muscle contraction (e.g. *Tnnt3* and *Atp2a1*) were reduced in both populations (Fig. 1e).

To get an overview of the NMJ-enriched transcripts, we performed Gene Ontology (GO)-term analysis. The results included terms such as basement membrane, neuromuscular junction, cell-cell junction, synapse and axon (Fig. 1f). This indicates that NMJ myonuclei express a range of transcripts that specifically support NMJ formation and maintenance. Interestingly, *Dok7* and *Rapsn,* encoding two essential proteins involved in forming and maintaining the NMJ ^29, 30^, were not significantly enriched in NMJ myonuclei, similar to what was observed with laser capture microdissection (LCM) ^31^. In conclusion, the method to isolate nuclei from mouse skeletal muscles without sorting them *via* FACS ^2, 3, 5^ or after genetic labeling ^4^ allowed us to identify eleven mono-nucleated cell populations, some of which were not previously reported. The method also yielded 472 and 501 transcripts that were highly enriched, and 270 and 379 transcripts that were reduced in NMJ and MTJ myonuclei, respectively. Interestingly, a good number of NMJ and MTJ transcripts overlap, suggesting molecular similarities between the two sub-compartments. Our data also show that transcripts encoding most (but not all) proteins essential for the formation and maintenance of the NMJ are specifically expressed in NMJ myonuclei.

### NMJ genes respond differently to denervation by sciatic nerve transection

The motor neuron orchestrates gene expression in myonuclei locally by secreting neural agrin, a signaling and scaffold-inducing protein, and by the release of acetylcholine, which triggers action potentials along the entire postsynaptic muscle fiber. Several lines of evidence indicate that neural agrin maintains synaptic gene expression at NMJ myonuclei through Lrp4/MuSK signaling and that electrical activity inhibits their expression in body myonuclei by calcium release into the cytoplasm ^20^. As a consequence, many NMJ transcripts, such as *Chrna1*, *Chrnb1*, *Chrnd*, ^10^ *Ncam1* ^32^ and *Musk* ^13^, are re-expressed (as during development) in body myonuclei upon denervation. This response is thought to be based on an ‘E-box’ domain that is bound by myogenic regulatory factors, such as myogenin, which are re-expressed upon electrical inactivity ^20^. However, not all NMJ genes contain such an ‘E-box’ domain within their promoter region. For example, *Chrne* transcripts remain largely restricted to NMJ myonuclei even after denervation ^10^.

To get a more global view on the regulation of NMJ genes, we next examined gene transcription using snRNA-seq from TA and GAS muscle after sciatic nerve transection. Bulk RNA-seq data show that TA and GAS muscles exhibit denervation gene signatures already three days post nerve transection while muscle atrophy takes seven days to become significant ^33, 34^. Hence, we chose 5-days post nerve transection as time point of analysis. The snRNA-seq data from denervated muscles were clustered together with the “innervated” dataset from Fig. 1. Similar to previous observations ^35, 36^, five-day denervation significantly increased the proportion of macrophages and Schwann cells compared to the innervated state (Supplementary Fig. S2a-c). FAP numbers have previously been shown to increase during denervation ^37, 38^, however at 5 days post denervation we observed no proportional increase with snRNA-seq. Interestingly, within the body myonuclei populations, two new clusters emerged (denervation 1 and 2), both expressing NMJ genes, such as *Musk* and *Ncam1* (Fig. 2a, b). Many fiber type-specific genes, such as *Mylk4* and *Myh4* were downregulated by denervation (Fig. 2b, Supplementary Fig. S2d). Denervation 1 and 2 were both devoid of *Myh4*, *Myh1* and *Myh2* transcripts (Supplementary Fig. S2d), which made it difficult to assign a specific fiber type to the clusters. Nevertheless, based on the localization of the UMAP clusters and other novel fiber type gene markers, such as *Agbl1* (Fig. 2b), nuclei in the denervation 1 cluster likely derive from type IIB fibers and nuclei in denervation 2 from IIA/IIX fibers. Furthermore, both denervation 1 and 2 clusters expressed distinct genes, such as *Scn5a* (denervation 1) and *Xirp2* (denervation 2; Fig. 2b), indicating that the denervation response differs between fiber types.

**Fig. 2:**
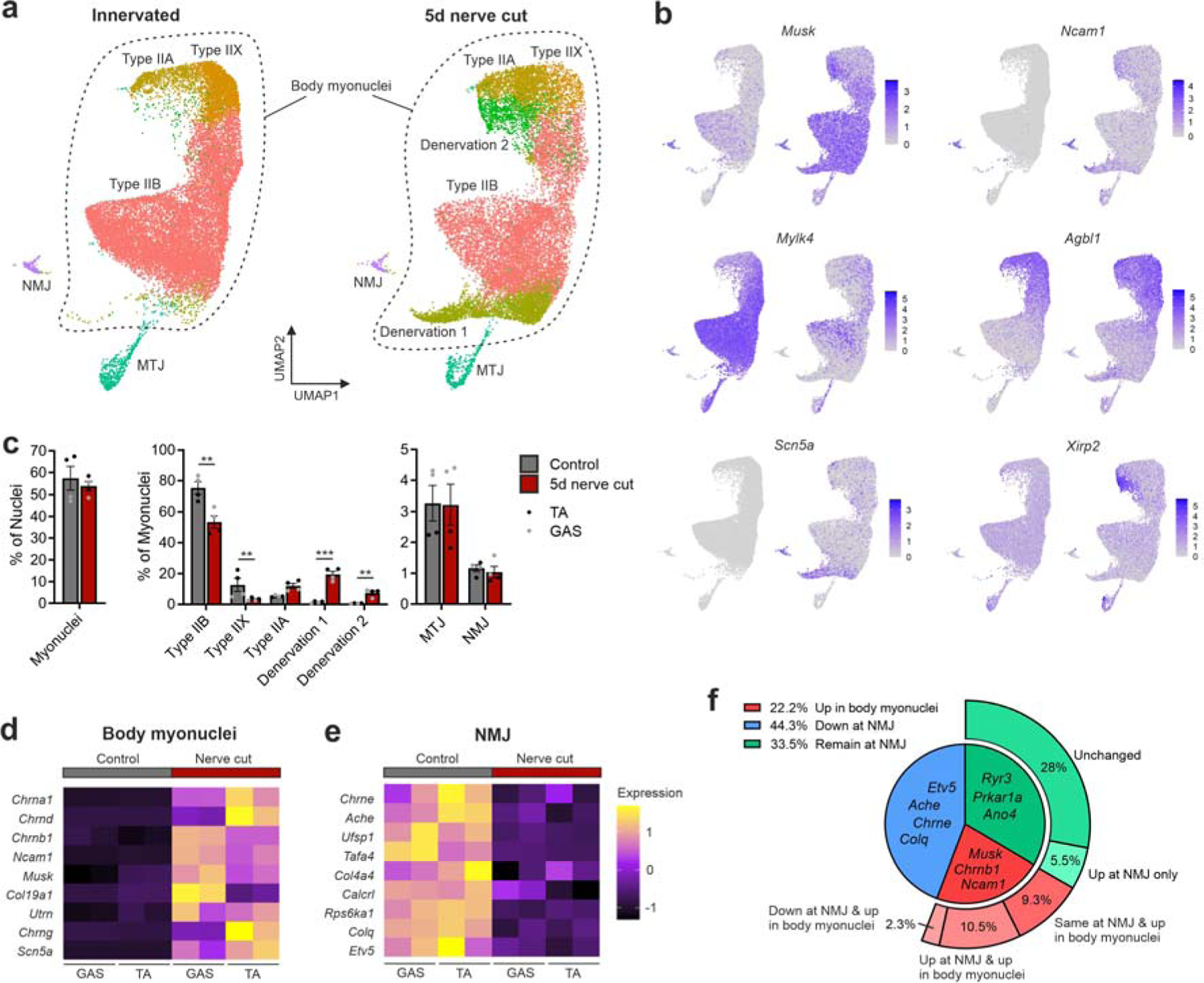
NMJ genes respond differently to denervation by sciatic nerve transection. **a** SnRNA-seq UMAP generated from control and five days denervated muscles. Only the myonuclei are visualized and the UMAP is split by condition to visualize differences in clustering. *n* = 2 for TA and GAS for both conditions. **b** Feature plots showing marker genes expressed in control and denervated myonuclei. **c** Proportion of myonuclei and effect of five days denervation. **d** Heatmap showing example genes upregulated in denervated body myonuclei. **e** Heatmap showing example genes downregulated in NMJ myonuclei after sciatic nerve cut. **f** Pie-chart visualizing the distribution of how NMJ transcripts (472 transcripts) change in expression after sciatic nerve cut in body and NMJ myonuclei. In (**c**) the data is presented as mean ± SEM and a regular t-test was performed (p < 0.05 = *, p < 0.01 = **, p < 0.001 = ***). In (**f**) transcripts were categorized as significantly changed when p-val. < 0.01 and log2FC ±0.25.

Denervation significantly affected the proportions within the myonuclei population (Fig. 2c). Despite observing a shift in major myonuclei populations in the UMAP after five days of denervation, NMJ and MTJ cluster sizes and localization were not significantly altered (Fig. 2a and Fig. 2c), suggesting that their transcriptional identities are largely independent of innervation.

Denervation resulted in presence of many NMJ transcripts, such as *Chrna1*, *Chrnb1, Chrnd, Musk* and *Ncam1* in body myonuclei (Fig. 2d), supporting the notion that denervation resets muscle fibers into a “developmental stage” as the transcriptional inhibition *via* electrical activity is removed ^10, 21, 32^. We observe the same pattern for other NMJ transcripts such as *Utrn* and *Col19a1*. In addition, denervation led to the re-expression of developmental genes, such as *Chrng* and *Scn5a*, that are silenced in innervated muscle (Fig. 2d). However, several prominent NMJ transcripts, such as *Chrne, Ache, Colq* and *Etv5* were not detected in body myonuclei and were even downregulated in NMJ myonuclei (Fig. 2e). By analyzing all NMJ-enriched transcripts (from Fig. 1), we found that ∼22% of the NMJ transcripts were upregulated in body myonuclei and hence may harbor an ‘E-box’ binding domain (Fig. 2f). However, almost half (∼44%) of the NMJ transcripts were not detected in body myonuclei upon denervation and were even downregulated in NMJ myonuclei. Another 28% of the NMJ transcripts were not significantly changed in their expression in NMJ myonuclei while ∼5% were specifically upregulated. These results suggest that either nerve-dependent “trophic” factors or the local depolarization (as a result of the release of ACh from the presynaptic nerve terminal) promote gene expression in NMJ myonuclei (Fig. 2f).

### Both trophic factors and electrical activity promote synaptic gene expression in NMJ myonuclei

While denervation by nerve transection removes the release of ACh from the nerve terminal and the suppressive function of electrical activity in body myonuclei, local agrin-Lrp4/MuSK signaling at the NMJ remains intact as agrin remains stably bound to the synaptic basal lamina ^39^ (Supplementary Fig. S3a). To identify the upstream pathways that regulate expression of particular NMJ genes, we next injected BoTX into TA muscles or blocked agrin-Lrp4/MuSK signaling at the NMJ by using the AAVMYO-CRISPR/Cas9 method to delete *Musk* ^40, 41^. BoTX blocks ACh release from nerve terminals and thereby blocks electrical activity in postsynaptic muscle fibers, but leaves the presynaptic nerve terminal intact ^42, 43^. Gene expression patterns were subsequently measured by snRNA-seq and the resulting data was clustered together with the dataset obtained from nerve transection shown in Fig. 2. The resulting UMAP similarly gave rise to denervation cluster 1, however denervation cluster 2 remained within the type IIA myonuclei cluster (Fig. 3a, Supplementary Fig. S3b-d). Five days after BoTX injection, myonuclei showed a denervation gene signature similar to nerve transection, with *Musk* and *Chrnb1* re-expressed in body and MTJ myonuclei (Fig. 3b). Three weeks post injection of AAVMYO ^41^ encoding *Musk*-single guide RNA (sgRNA) into the TA, *Musk* transcripts were strongly depleted at the NMJ (Fig. 3b) and AChR density, visualized by whole mount staining, was reduced (Fig. S3e). Nevertheless, *Chrnb1* was not re-expressed in body myonuclei (Fig. 3b), confirming acute MuSK depletion was yet to induce secondary signaling effects related to denervation, as seen with chronic MuSK depletion ^40^. *Musk* knockout (KO) significantly downregulated a discrete set of genes, not downregulated by nerve transection or BoTX injection, including *Prkar1a* and *Ryr3* (Fig. 3b). This shows that the reduction of MuSK protein (i.e. agrin signaling) was sufficient to affect a component of NMJ gene expression distinct from both innervation and denervation.

**Fig. 3:**
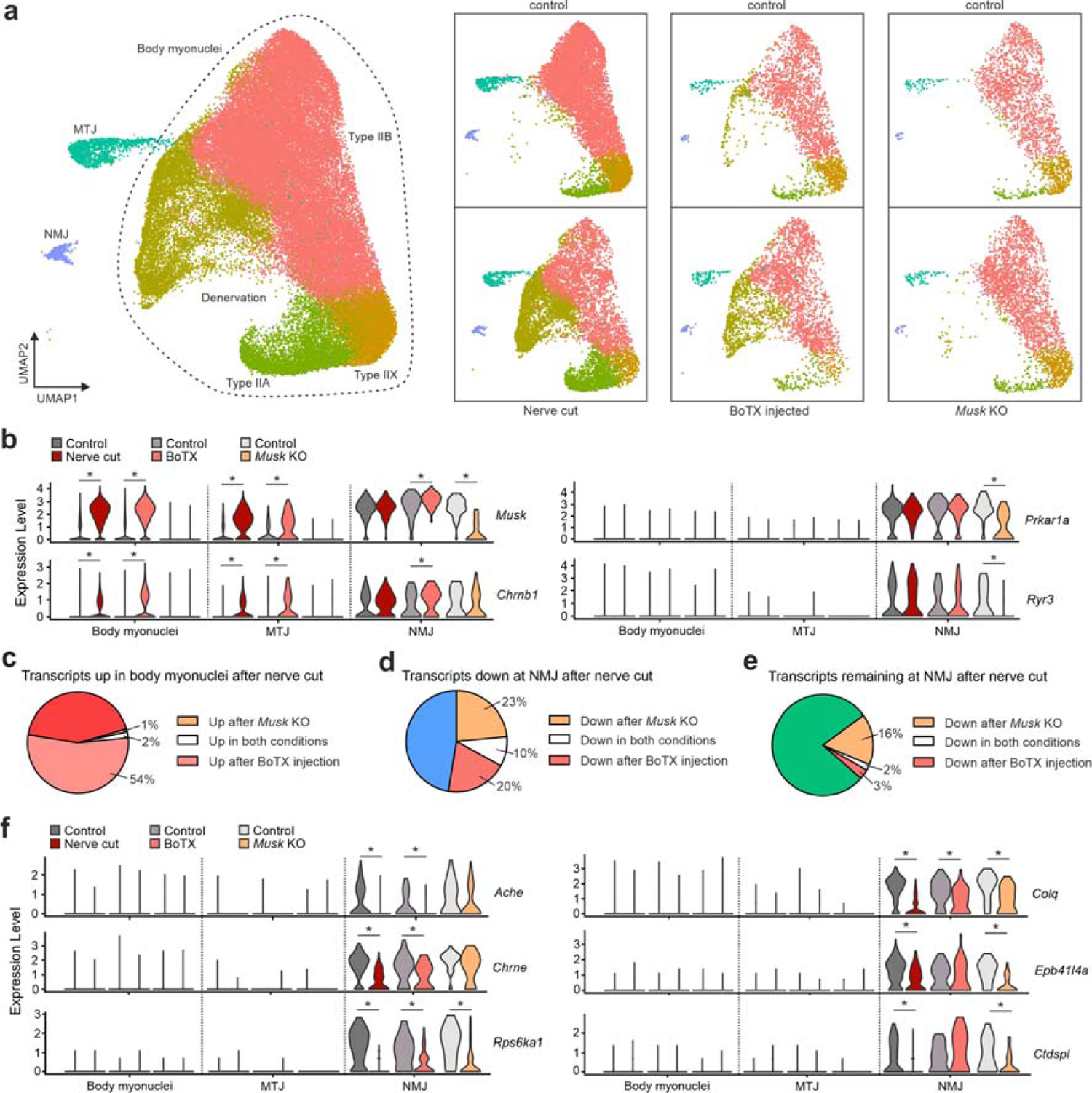
Both trophic factors and electrical activity promote synaptic gene expression in NMJ myonuclei. **a** SnRNA-seq was performed on BoTX injected and *Musk* knockout (KO) TA muscles and respective controls and the data were clustered together with the previous nerve cut dataset from Fig. 2. Only myonuclei (∼60’000) are depicted in the UMAP. Individual conditions and their controls are visualized as split UMAPs to the right. **b** Violin plots visualizing the expression of selected NMJ genes in myonuclei after nerve cut, BoTX injection and *Musk* KO. The violin plots were generated from snRNA-seq data, UMAP is depicted in (**a**). Body myonuclei from all fiber types were pooled together to generate an average expression value. **c**, **d** and **e** Pie charts showing how synaptic genes are regulated after BoTX injection and *Musk* KO compared to sciatic nerve transection. **f** Violin plots of NMJ genes showing different responses to nerve cut, BoTX injection and *Musk* KO, indicating different gene regulatory mechanisms. Nerve cut and the corresponding controls *n* = 4, the remaining conditions *n* = 1. All transcripts with a p-val. < 0.01 and log2FC ±0.25 are indicated by *. In (**c**-**e**) transcripts were categorized as significantly changed when p-val. < 0.01 and log2FC ±0.25.

After confirming successful denervation by BoTX injection and reduction of agrin-Lrp4/MuSK-signaling, we explored NMJ gene expression in body and NMJ myonuclei. Out of all the NMJ genes re-expressed in body myonuclei by sciatic nerve transection, 55% were also re-expressed by BoTX (Fig. 3c). In contrast, only very few transcripts were re-expressed in MuSK-depleted muscle (Fig. 3c), suggesting that the muscle fibers were not yet denervated. However, in NMJ myonuclei both BoTX injection and MuSK depletion resulted in the downregulation of many NMJ-enriched transcripts (Supplementary Fig. S3f). Interestingly, out of the 218 transcripts that were downregulated in NMJ myonuclei after MuSK depletion, 123 (56%) were NMJ transcripts (Supplementary Fig. S3f), suggesting that lack of agrin-MuSK signaling is a major contributor to the maintenance of local transcription at the NMJ.

To assess the contribution of factors that require the presence of the presynaptic nerve but are independent of agrin-Lrp4/MuSK signaling, we next examined the overlap of transcripts in NMJ myonuclei that were downregulated by nerve cut (loss of nerve and electrical activity) with BoTX injection (preservation of presynaptic nerve and loss of electrical activity) and MuSK depletion (preservation of presynaptic nerve and electrical activity but reduction of agrin-Lrp4/MuSK signaling). When comparing the downregulated genes after nerve transection with the downregulated genes after BoTX injection and *Musk* knockout, ∼30% and ∼33% respectively showed the same pattern (Fig. 3d). As we observed downregulation of NMJ genes after both nerve cut and BoTX injection, nerve-evoked electrical activity seems to promote the expression of certain NMJ genes such as *Ache* and *Chrne* (Fig. 3f). Several transcripts such as *Rps6ka1* and *Colq* were reduced in all conditions, which indicates that certain genes need a combination of MuSK signaling and electrical activity (Fig. 3f). Since we observed downregulation of certain NMJ genes by nerve cut and *Musk* KO but not by BoTX injection (*Ctdspl* and *Epb41l4a* matching this criteria, Fig. 3f), this supports the hypothesis that other potential trophic factors, e.g., neuregulins ^20^, play a role in promoting local gene expression at the NMJ. Next, we quantified the contribution of agrin-Lrp4/MuSK signaling during denervation and hence overlapped NMJ transcripts that remain at the NMJ after nerve cut to BoTX injection and *Musk* KO. As expected, BoTX injection showed a very similar response to nerve cut (Fig. 3e). *Musk* KO lead to a reduction of 18% of these “denervation resistant” transcripts (Fig. 3e). This confirms that agrin persisting in the synaptic basal lamina after denervation is sufficient to promote expression of certain NMJ genes.

In conclusion, we confirm that agrin-Lrp4/MuSK signaling plays a critical role in synaptic gene expression at the NMJ. Interestingly, our data suggest that depolarization as well as trophic factors, other than agrin, contribute significantly to gene expression in NMJ myonuclei (Supplementary Fig. S3g).

### Selection and confirmation of NMJ genes

Similar to other snRNA-seq datasets ^2, 3, 4^, we detected numerous uncharacterized NMJ-specific and - enriched transcripts. As we were interested in better understanding mechanisms involved in NMJ maintenance, we selected transcripts that were specific to NMJ myonuclei in innervated muscle, that were not re-expressed by body myonuclei in denervation and whose expression was downregulated by *Musk* KO (Supplementary Fig. S4a, b). We ultimately selected three genes, *Lrtm1, Pdzrn4* and *Etv4* for a detailed functional analysis based on their expression pattern and some additional functional criteria. *Lrtm1*, that was upregulated after sciatic nerve transection and BoTX injection specifically in NMJ myonuclei (Supplementary Fig. S4a) encodes a member of the leucine-rich repeat transmembrane protein family, which have been implicated in neural development ^44^. The second candidate was *Pdzrn4* (Supplementary Fig. S4a), which encodes a PDZ domain-containing protein paralogous to PDZRN3 that is expressed in body and NMJ myonuclei and has been shown to bind to MuSK and target it for degradation in cultured myotubes ^45^. Pdzrn4 was downregulated after sciatic nerve transection and MuSK KO but not after BoTX injection indicating that it may be partially regulated by a trophic factor other than agrin. Finally, we selected *Etv4*, a member of the ETS family of transcription factors that have been implicated in the regulation of NMJ gene expression by their binding to the ‘N-box’ domain ^20^. Indeed, the global knockout of *Etv5* has been show to affect the NMJ^46^. Interestingly, *Etv4*, similar to *Etv5*, showed a tendency to be downregulated in all our experimental paradigms (Supplementary Fig. S4a) indicating that they are regulated by the same factors.

First, we confirmed expression of the transcripts near NMJ myonuclei by using smFISH. *Etv4*, *Lrtm1* and *Pdzrn4* transcripts were highly enriched near NMJ myonuclei (identified by probes against *Chrne*) of both, fast-twitch TA and slow-twitch *soleus* muscles (Fig. 4a, b). All three transcripts were also enriched in NMJ regions isolated from TA muscle by laser capture microdissection ^31^ (Fig. 4c).

**Fig. 4:**
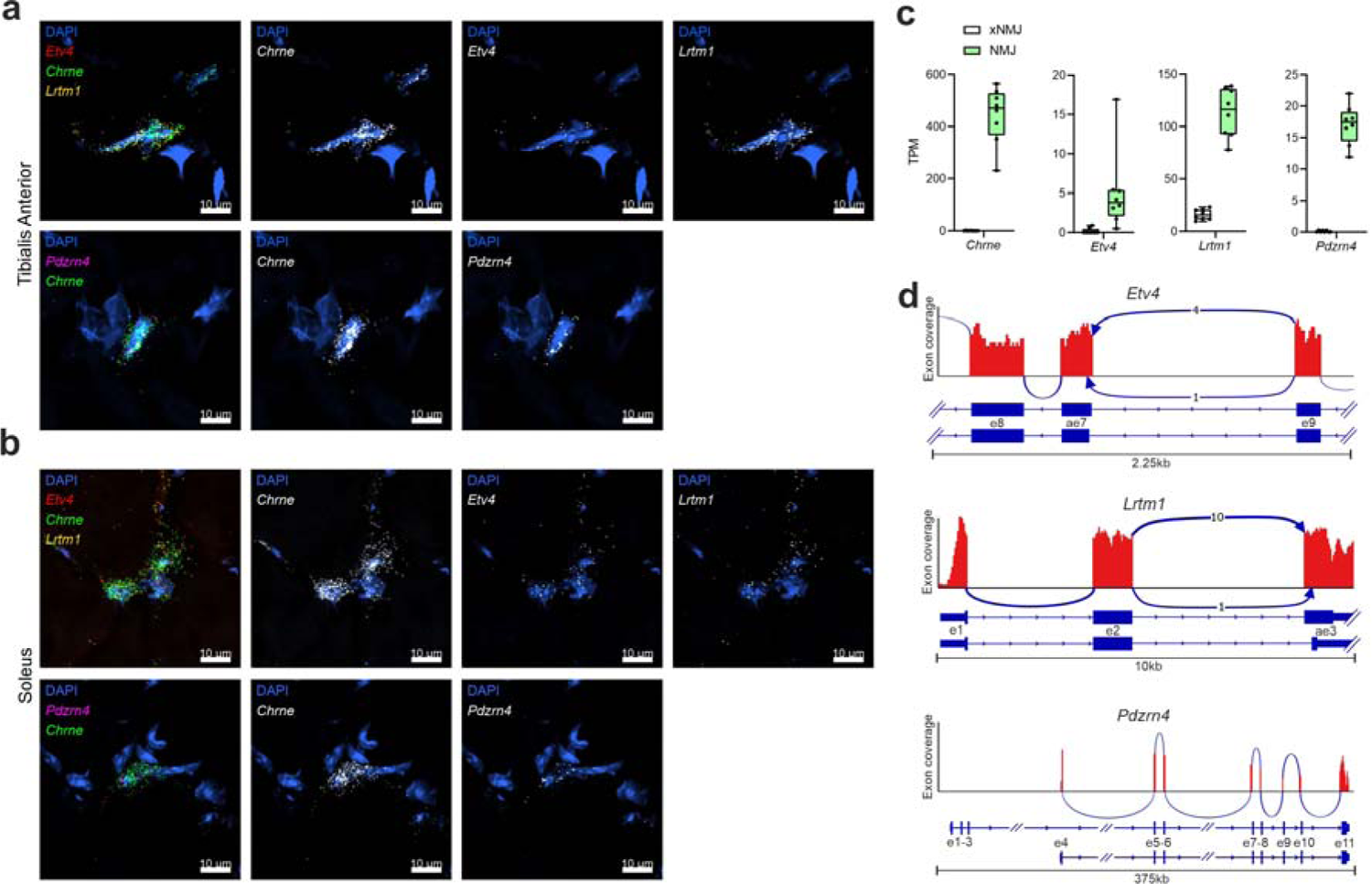
Selection and confirmation of NMJ genes. **a** and **b** single molecule fluorescent *in situ* hybridization (smFISH) on TA and *soleus* cross-sections from adult C57BL/6 mice. NMJ myonuclei were identified by the presence of *Chrne* transcript. **c** Box-plots showing average transcripts per million (TPM) values from laser-capture-isolated NMJ regions and non-NMJ regions (xNMJ) ^31^. **d** Representative Sashimi plots generated from bulk-RNA-seq data of TA and *soleus* ^31^. For *Pdzrn4* and *Lrtm1*, all exons (e) are depicted and for *Etv4*, only three exons are depicted (a region that undergoes alternative mRNA splicing). Exons are portrayed as blue boxes with the aligned reads in red. The numbers in-between splice junctions show the proportion of how frequently a particular overlapping read was detected. Exons that can be spliced alternatively at their 3’ or 5’ end are indicated as alternative exons (ae).

To assure that we would study the correct protein isoform of the three candidate genes, we used bulk RNA-seq data for TA and *soleus* ^31^ to determine the predominate splice variants expressed in skeletal muscle (Fig. 4d, Supplementary Fig. S4c-e). Reads mapping to exon junctions showed the existence of different transcripts for *Etv4* and *Lrtm1*. Interestingly, *Pdzrn4* encodes two different transcripts with alternative transcription start sites, only one of which was found in muscle (Fig. 4d, Supplementary Fig. S4d). This transcript codes for a protein that lacks the N-terminal RING domain and hence makes it unlikely that PDZRN4 acts as an E3 ubiquitin ligase as was described for PDZRN3 ^45^. In conclusion, the NMJ transcripts *Etv4* and *Lrtm1* undergo alternative mRNA splicing with a preponderance of one particular splice variant and for *Pdzrn4* only one transcript version was found.

### PDZRN4 overexpression mislocalizes NMJ proteins and ETV4 is sufficient to induce NMJ transcripts in body myonuclei

To characterize protein localization and function of the candidate genes in mouse skeletal muscle, we next overexpressed the most dominantly expressed muscle splice versions of ETV4, LRTM1 and PDZRN4 by intra-muscular injection of myotropic AAVMYO ^41^ into the TA (Fig. 5a). Expression was driven by the CMV promoter and proteins were FLAG-tagged at their C-terminal end (Fig. 5a). Lower hindlimb muscles were collected one and two weeks post injection. The overexpressed proteins were localized by IF staining on whole-mount preparations (Fig 5b) and cross-sections (Supplementary Fig. S5a). All proteins were enriched at the NMJ as visualized by AChR labeling with α-bungarotoxin (Fig. 5b). The transcription factor ETV4 localized to myonuclei and LRTM1 to the plasma membrane (Fig. 5b, Supplementary Fig. S5a, b). PDZRN4 formed punctate stripes within muscle fibers (Fig. 5b), presenting as vesicle-like structures in cross-sections (Supplementary Fig. S5a). PDZRN4 also promoted and was associated with extrasynaptically located AChR and MuSK clusters (Fig. 5c, Supplementary Fig. S5c). Although PDZRN4 could be visualized throughout the entire muscle fiber, the MuSK- and AChR-positive puncta remained in close proximity of the NMJ, suggesting AChR and MuSK gene transcription and protein translation remains specific to NMJ myonuclei.

**Fig. 5:**
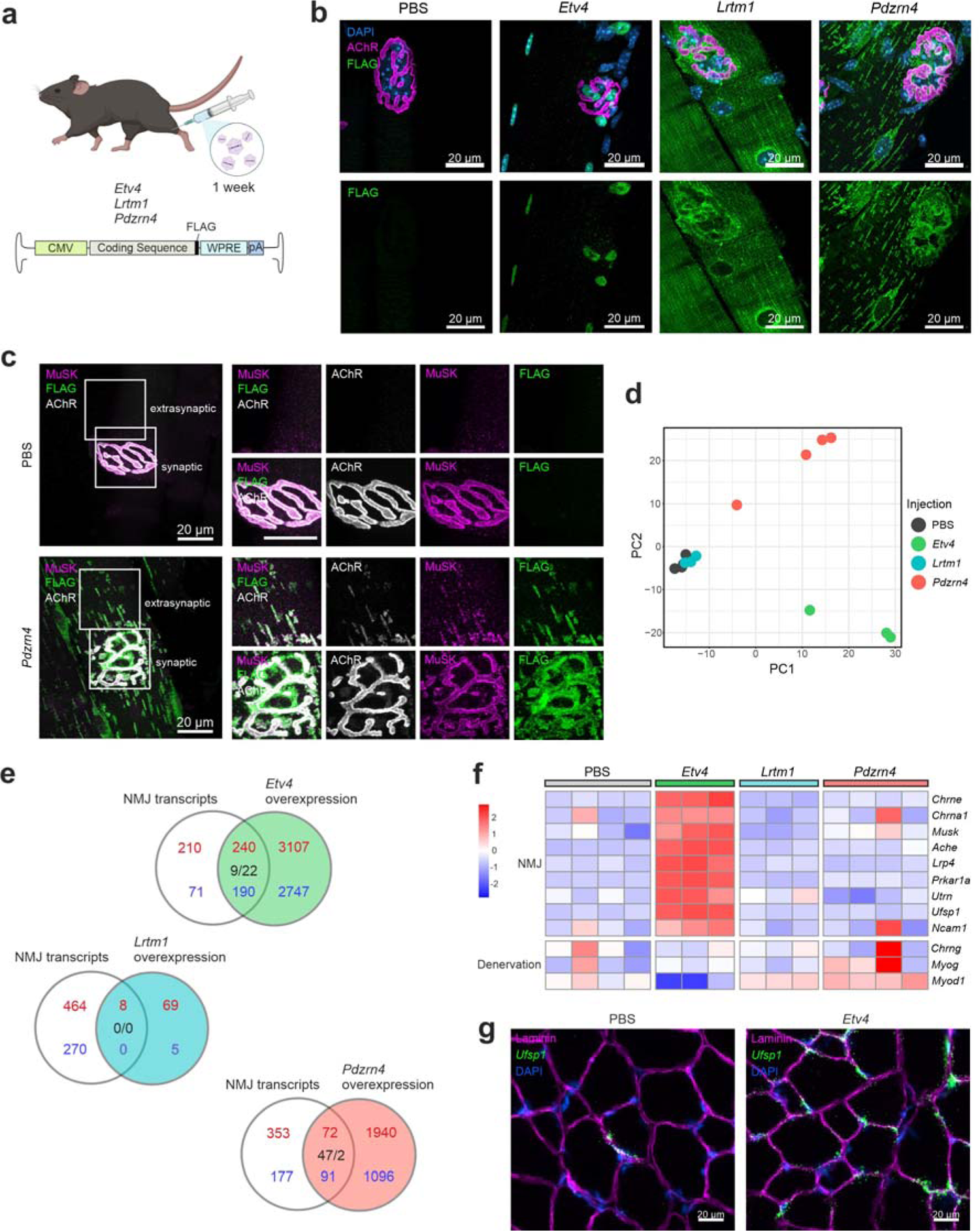
PDZRN4 overexpression mislocalizes NMJ proteins and ETV4 is sufficient to induce NMJ transcripts in body myonuclei. **a** Experimental scheme; AAVMYO containing either *Etv4*, *Lrtm1* or *Pdzrn4* coding sequence was injected into the TA of wild-type adult mice (2.5 x 10^10^ vector genome copies). All three targets were driven by a CMV promoter and were FLAG-tagged at the C-terminal end. **b** One-week post injection, whole-mount preparations of the *extensor digitorum longus* (EDL) were stained with DAPI, α-bungarotoxin and an anti-FLAG antibody. **c** Whole-mount preparations of *Pdzrn4*- and PBS-injected EDL muscles. Overexpression of PDZRN4 leads to extrasynaptic localization of AChRs and MuSK. **d** Principle component (PC) analysis of bulk RNA-seq data from TA muscles overexpressing either *Etv4*, *Lrtm1* or *Pdzrn4*. **e** Venn diagrams comparing NMJ transcripts (from Fig. 1) with up- or downregulated transcripts induced by overexpression of the indicated gene. Numbers in red/blue indicate NMJ enriched/depleted transcripts or upregulated/downregulated genes, respectively. Numbers in black represent oppositely regulated genes. **f** Heatmap generated from bulk RNA-seq depicting how NMJ and denervation-associated genes are changed by overexpression. **g** SmFISH of the NMJ transcript *Ufsp1* ^3^ combined with an IF staining for laminin on TA cross-sections. Experimental scheme in (**a**) was created with BioRender.com. In (**e**) up- and down-regulated transcripts fulfill padj. < 0.05 and Log2FC ± 0.25.

Next, we analyzed the muscle transcriptome using bulk-RNA-seq in all samples. According to principle component (PC) analysis and the number of differentially expressed genes, both ETV4 and PDZRN4 strongly affected the muscle transcriptome, while LRTM1 had little effect (Fig. 5d, Supplementary Fig. S5d). Transcriptional changes were greatest with ETV4, increasing expression of more than 3,000 transcripts and decreasing a similar number, followed by PDZRN4, which increased expression of 2’000 transcripts and reduced more than 1,000 (Supplementary Fig. S5d). To get an idea as to how the NMJ might be affected, we compared the list of up- and downregulated genes to that of the NMJ transcripts (from Fig. 1). Strikingly, overexpression of ETV4 resulted in increased expression of more than 50% of the NMJ transcripts (Fig. 5e). These transcripts included well-characterized transcripts, such as *Chrne*, *Chrna1*, *Musk*, *Ache* and *Lrp4* (Fig. 5f). Importantly, the upregulation of NMJ genes in ETV4-overexpressing muscles was not a consequence of denervation, as denervation associated genes such as *Chrng*, *Myog* and *Myod1* ^34, 47^ were not upregulated (Fig. 5f). In contrast, LRTM1 and PDZRN4 increased only 1.7% and 15% of the NMJ transcripts, respectively (Fig. 5e) and none of the well-known NMJ transcripts were increased in these two samples (Fig. 5f). To finally test, whether ETV4 increased the expression of NMJ transcripts in all myonuclei or only in NMJ myonuclei, we used smFISH against the NMJ transcript *Ufsp1* ^3^. While *Ufsp1* was expressed at few sites in WT muscle, consistent with NMJ-restricted expression, muscle-wide ETV4 expression caused widespread *Ufsp1* expression, with RNA puncta detected in most body myonuclei (Fig. 5g). All in all, these experiment show that forced ETV4 expression is sufficient to potently stimulate an NMJ-specific transcriptional program in body myonuclei, in line with the notion that a large portion of NMJ transcripts are regulated by ETS transcription factors ^46^.

As the top DAVID GO-term in the bulk RNA-seq samples from muscles overexpressing either ETV4 or PDZRN4, but not LRTM1, was “immune system process” (Supplementary Fig. S5e), we asked whether prolonged expression would affect muscle health. Indeed, muscle fibers overexpressing PDZRN4 or ETV4 for two weeks showed signs of muscle degeneration and regeneration, indicated by the presence of mouse IgG inside of the muscle fibers and of many centrally located myonuclei (Supplementary Fig. S5f). These results indicate that long-term ectopic expression of proteins that affect the expression (ETV4) or localization (PDZRN4) of proteins confined to the NMJ (i.e. MuSK and AChR) is not well tolerated. The fact that LRTM1 overexpression does not result in such a muscle response shows that the pathological changes are not caused by the overexpression of any protein or are a response to AAVMYO transduction.

### Muscle fiber-specific knockout of Pdzrn4 causes NMJ fragmentation

While overexpression studies test for sufficiency of a protein, addressing essentiality requires their removal from the organism. To see whether any of the candidates were required to maintain NMJ function and structure, we used our recently established muscle fiber-specific AAV-CRISPR/Cas9 system ^40^ and designed six to seven sgRNAs for each gene (Supplementary Fig. S6a). Upon cloning them into an AAV vector, we delivered them using AAVMYO ^41^ to mice expressing Cas9 in skeletal muscle fibers (Cas9mKI). We injected 3 x 10^11^ sgRNA-encoding AAVMYO (∼9 x 10^12^ vector genomes/kg) *via* the tail vein into Cas9mKI mice to generate systemic muscle fiber-specific knockouts (Fig. 6a). Fifteen weeks post injection, mice were analyzed and transcript levels for the targeted genes were measured in TA by qPCR. In all cases, targeted transcripts were significantly lower than in control TA (Fig. 6b). Interestingly, *Musk* mRNA was also significantly downregulated by knockout of *Etv4, Lrtm1* and *Pdzrn4* (Fig. 6b; right), suggesting an important contribution of all genes to synaptic signaling. The extent of transcript loss differed between the targeted genes, which can be due to different extent of nonsense-mediated mRNA decay and mRNA expression in muscle-resident cells other than muscle fibers. For example, *Etv4* is also expressed by tenocytes (Supplementary Fig. S4b). Functionality of the neuromuscular system was measured using all-limb and forelimb grip strength as well as balance beam performance 10-weeks post injection. None of these measures differed between the knockout and controls (Fig. 6c, Supplementary Fig. S6b). Similarly, no significant decrement in action potential was observed after 10 stimulations at 40 Hz frequency using compound muscle action potential (CMAP) measurements 15-weeks post sgRNA injection (Fig. 6d). In conclusion, knockout of *Etv4*, *Lrtm1* or *Pdzrn4* in adult skeletal muscle fibers does not result in a significant deficit in neuromuscular performance 15-weeks post injection. It is important to note that the NMJ is characterized by its high safety factor, which assures reliable generation of an action potential even at low AChR density ^48^. Hence, NMJs could still be affected at the molecular level without functional consequences at this 15-week time point. To investigate whether morphological changes could be observed, we performed whole-mount IF stainings of the *extensor digitorum longus* (EDL). IF imaging revealed that both *Etv4* and *Lrtm1* knockout mice showed non-significant trends towards a higher number of fragments (Fig. 6e, f). In *Pdzrn4* knockout mice, the number almost doubled and reached high significance (Fig. 6g). As NMJ function was seemingly not altered in any of the knockouts, it remains an open question whether NMJ fragmentation could be an early sign for its instability. It is well known that NMJ fragmentation in rodents is a hallmark of aging although this structural change also does not result in functional impairments ^49^.

**Fig. 6:**
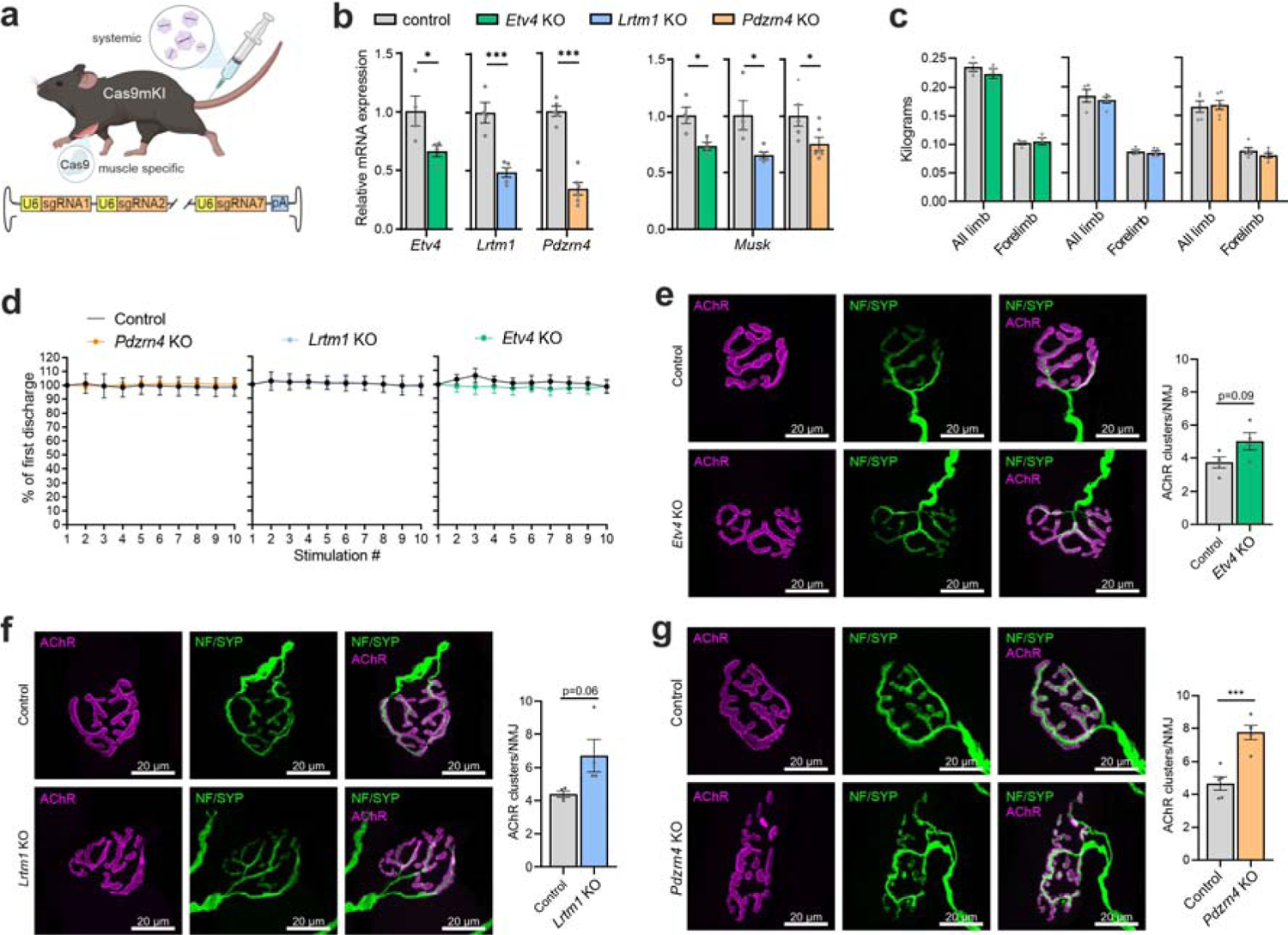
Muscle fiber-specific knockout of Pdzrn4 causes NMJ fragmentation. **a** Cas9mKI mice were injected with AAVMYO encoding 6 (*Lrtm1*) or 7 (*Etv4* and *Pdzrn4*) different sgRNAs (3 x 10^11^ vector genome copies) *via* the tail vein. **b** Relative mRNA expression of the targeted genes in TA muscle 15 weeks post-injection. Note that *Musk* mRNA was also significantly reduced in all three groups. **c** All limb and forelimb grip strength measurements were performed on mice 10 weeks post injection. **d** CMAP was performed on the GAS 15 weeks post injection at different frequencies (image shows average values at 40 Hz frequency). **e**, **f**, and **g** representative whole mount images of NMJs from the EDL from *Etv4* (**e**), *Lrtm1* (**f**) and *Pdzrn4* (**g**) knockout mice and control littermates. AChRs are visualized by α-bungarotoxin and the presynapse by neurofilament (NF) and synaptophysin (SYP). Bar graphs next to the images show the quantification of AChR fragmentation. NMJs of *Pdzrn4* knockout mice are significantly fragmented. Experimental scheme in (**a**) was created with BioRender.com. The data in (**b**-**g**) is presented as mean ± SEM. In (**f**-**g**) a two-tailed t-test was performed. p < 0.05 = *, p < 0.01 = **, p < 0.001 = ***. For whole mount quantifications, a minimum of 10 NMJs per mouse were quantified (*n* = 4 for *Etv4* and *Lrtm1* and *n* = 5 for *Pdzrn4* knockout groups).

### PDZRN4 localizes to the Golgi apparatus and binds MuSK

As PDZRN4 overexpression caused AChR and MuSK mislocalization and the muscle-specific knockout caused NMJ fragmentation, we focused on elucidating the function of PDZRN4 at the NMJ. IF stainings showed PDZRN4 localizing to vesicle-like structures (Fig. 5b, Supplementary Fig. S5a, c). As vesicles are transported across the microtubule network, we performed an IF staining against alpha-tubulin together with PDZRN4. FLAG-tagged PDZRN4 showed strong colocalization with microtubules (Fig. 7a), strengthening the hypothesis that these structure are indeed vesicles. To investigate which type of vesicle or organelle PDZRN4 associates with, PDZRN4 was overexpressed in HeLa cells, where we observed strong colocalization with the Golgi apparatus (Fig. 7b). Interestingly, in muscle fibers the Golgi apparatus is not only in close proximity to nuclei (as in other cells), but also extends throughout the muscle fiber *via* the microtubule network ^50, 51, 52, 53^, as we observed for PDZRN4.

**Fig. 7:**
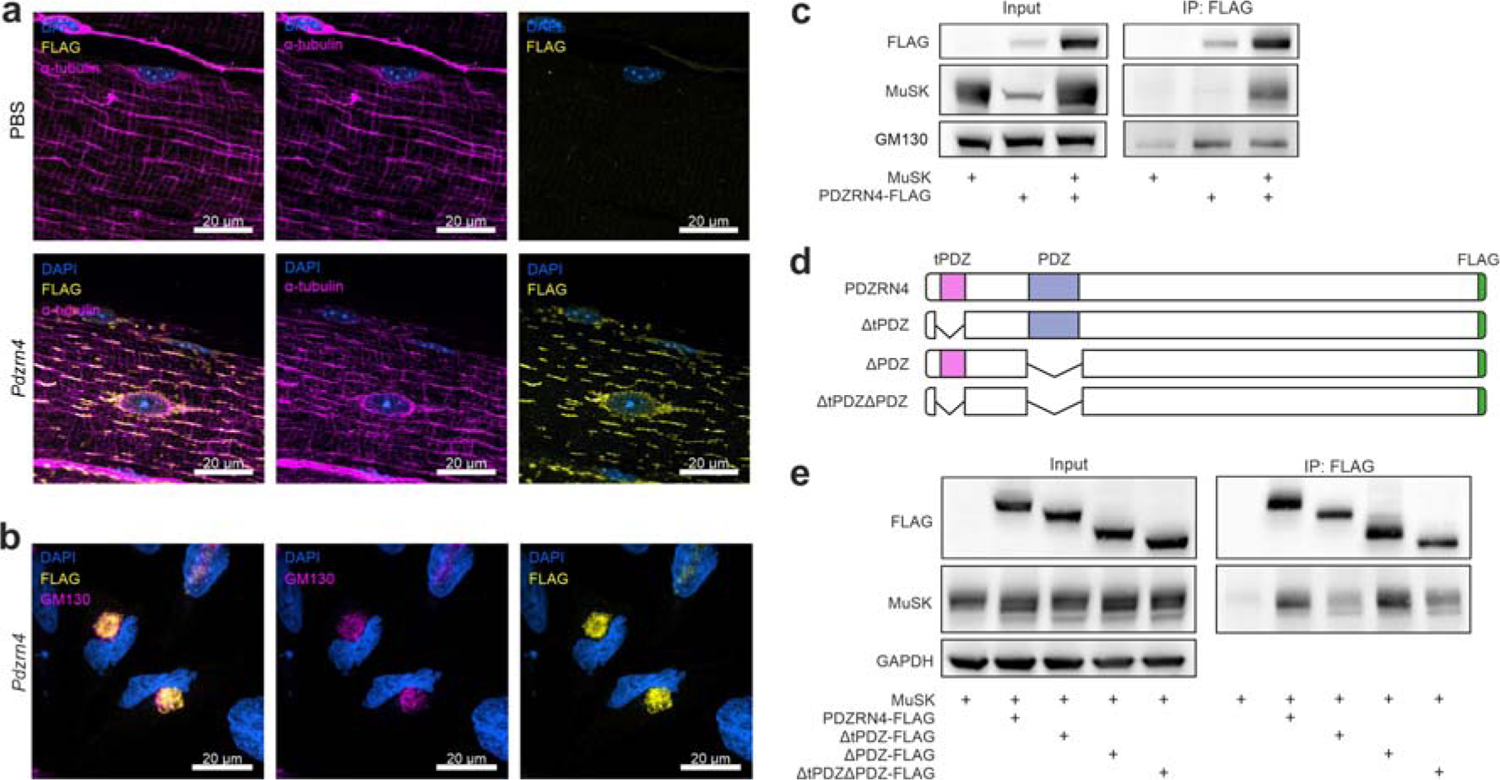
PDZRN4 localizes to the Golgi complex and binds MuSK. **a** IF stainings of whole mount prepared EDLs show that PDZRN4 strongly colocalizes with microtubules. **b** HeLa cells were transfected with the same *Pdzrn4* plasmid used in mouse experiments (depicted in Fig. 5a). PDZRN4 colocalized with the Golgi marker GM130. **c** Immunoprecipitation (IP) was conducted in HEK 293T cells overexpressing PDZRN4 and/or MuSK. IP against the FLAG tag of PDZRN4 resulted in co-pull down of both MuSK and GM130. **d** The PDZRN4 isoform expressed in muscle contains a conventional and a truncated PDZ domain (tPDZ). Each domain was removed to investigate their respective contributions to MuSK interaction. **e** IP experiment using the deletion mutants of PDZRN4 shown in (**d**). The interaction between MuSK and PDZRN4 is independent of the conventional PDZ domain.

As PDZRN4 overexpression in skeletal muscle affects MuSK and AChR localization, we next tested whether PDZRN4 would co-immunoprecipitate (IP) with MuSK. Indeed, when MuSK and FLAG-tagged PDZRN4 were both overexpressed in HEK 293T cells, MuSK was co-enriched in the anti-FLAG IP (Fig. 7c). Interestingly, GM130 was also detected in the IP, indicating that some Golgi vesicles remain intact upon the mild lysis used for this experiment (Fig. 7c). As PDZRN4 associates with MuSK and the Golgi apparatus, we co-overexpressed both proteins in HeLa cells to see how protein localization would be affected. IF staining revealed intracellular MuSK accumulation together with PDZRN4 in a Golgi-like pattern (Supplementary Fig. S7a).

To map the binding site of PDZRN4 to MuSK we next generated deletion mutants. The *Pdzrn4* transcript expressed in skeletal muscle encodes one conventional PDZ domain as well as a truncated PDZ domain (called herein tPDZ), close to its N-terminus (Fig. 7d, Supplementary Fig. S7b). MuSK binding in the PDZRN3 paralog has been mapped to its PDZ domain ^45^. To test whether this would also be the case for PDZRN4, the PDZ and the tPDZ domain were removed individually as well as in combination. Interestingly, while removal of the conventional PDZ domain did not affect co-IP of MuSK, removal of the tPDZ domain and of both domains strongly lowered the amount of MuSK (Fig. 7e). Although MuSK still appeared in the immunoprecipitate (Fig. 7e), deletion of the tPDZ domain strongly affected PDZRN4 association with the Golgi apparatus as observed in IF stainings in HeLa cells (Supplementary Fig. S7c). To test if removal of any other region in PDZRN4 would affect MuSK association, five additional regions of PDZRN4 were removed individually (Supplementary Fig. S7d). Co-IP depicted that all PDZRN4-FLAG versions were able to pull down MuSK, though the N-terminal region appeared to be of higher importance (Supplementary Fig. S7e).

In conclusion, these results indicate that PDZRN4 associates with MuSK at the Golgi apparatus. Furthermore, our data show that PDZRN4 does not require its PDZ domain to interact with MuSK, setting it apart from PDZRN3 ^45^.

## Discussion

NMJ myonuclei are a small population of nuclei that share one cytoplasm with hundreds of other myonuclei. Research over several decades has shown that these NMJ myonuclei express genes whose protein products aggregate at the postsynaptic side of the NMJ. Among those are essential NMJ components, such as Lrp4, MuSK and the AChR subunits. There are notable exceptions such as *Rapsn* and *Dok7*, whose protein products cluster at the NMJ ^30, 54^, but its transcripts are not more expressed by NMJ myonuclei compared to body myonuclei. While many NMJ genes are transcribed in body myonuclei during development, the formation of the NMJ by the clustering of these proteins at the site of nerve-muscle contact and the onset of nerve-evoked electrical activity, causes the NMJ myonuclei to become specialized to transcribe synaptic genes ^6, 20^. The continuous expression of these NMJ-essential genes in NMJ myonuclei throughout life suggests the presence of nerve-derived, local signals that maintain their transcription. In addition, the nerve-evoked electrical signal, which spreads within the muscle fiber, suppresses the expression of the synaptic genes in body myonuclei^20^. Bulk RNA-seq based methods using for example microdissection of synaptic and non-synaptic regions, do not allow to distinguish between transcripts from NMJ nuclei and those from other cell types localized at synaptic regions; e.g., myelinating and terminal Schwann cells ^31^. To overcome this challenge, we used snRNA-seq in combination with experimental manipulations of the innervation state to dissect the signaling pathways involved in synaptic gene expression.

We identified 472 transcripts that were significantly enriched in NMJ myonuclei. These included the well-characterized NMJ-associated genes *Musk, Lrp4, Ache, Colq, Ncam1, Etv5* and the different *Chrn* subunits, which validated our approach. Some of the transcript were enriched in both NMJ and MTJ nuclei (Fig. 1), suggesting structural and functional similarities between the two specializations. For example, transcripts for laminin-α2 (*Lama2*) and collagen IV (*Col4a1* and *Col4a2*) were enriched in both nuclei populations. Laminin-α2 is the major laminin chain and collagen IV the main collagen of the basement membrane surrounding skeletal muscle fibers ^55^. Interestingly, mutations in *LAMA2* cause a severe, early onset congenital muscular dystrophy ^56^. The fact that transcripts for these proteins are enriched in NMJ and MTJ nuclei indicates that there is an increased requirement of basement membrane in these specialized regions. In contrast, other members of the laminin and collagen protein family, such as *Lamb2*, *Col13a1* and *Colq*, were selectively transcribed in NMJ nuclei. Interestingly, mutations in all three genes cause congenital myasthenic syndrome ^57^. Similarly, *Col22a1* is selectively expressed in MTJ nuclei and the protein is well-known to be a specific marker for MTJs ^58^.

To get deeper insights into the regulation of synaptic gene expression in NMJ myonuclei, we also generated snRNA-seq data from denervated TA and GAS muscles five days after sciatic nerve transection (Fig. 2) and after NMJ perturbations *via* injection of BoTX (Fig. 3), which prevents the release of ACh from the presynaptic nerve terminal and thus causes chemical denervation without the physical removal of the nerve. This perturbation hence still allows the nerve to provide “trophic support” to the postsynaptic region. Finally, we also perturbed agrin-Lrp4/MuSK by depleting MuSK to abolish the signaling pathway essential for the formation and the maintenance of the NMJ ^6^. Denervation by nerve transection resulted in the loss of synapse-specific expression of a subset of genes; mainly based on the upregulation of NMJ genes in body myonuclei (Fig. 2). This is well described and mechanistically has been explained by the loss of nerve-evoked electrical activity that allows the re-expression of myogenic regulatory factors, such as myogenin and myoD ^47, 59, 60^. Unlike the NMJ-specific transcription factor ETV5, which binds to the ‘N-box’, myogenic regulatory factors bind to the ‘E-box’ domain and thereby promote the expression of NMJ genes ^20^. However, we observed that a large proportion of synaptic transcripts remained specifically expressed in NMJ myonuclei and some were even downregulated during denervation, suggesting that they do not harbor an ‘E-box’ domain. We therefore theorized that the downregulation of synaptic genes could be from either the loss of trophic factors provided by the nerve or by the loss of electrical activity. To test this, snRNA-seq was performed on chemically denervated muscle by BoTX injection, where the motor nerve terminal remains at the NMJ ^42, 43^. Approximately 30% of the NMJ genes downregulated by nerve cut were also downregulated by BoTX (Fig. 3), indicating that nerve-evoked electrical activity plays an important role in maintaining synaptic gene expression. A DNA binding protein that shows reduced nuclear localization during denervation, and therefore might be implicated in the reduced expression of synaptic genes, is the protein FUS. FUS has been proposed to regulate synaptic genes together with ETV5 and mutations of *Fus* can cause an aggressive form of ALS ^61^. Furthermore, we observed that certain NMJ transcripts were “resistant” to denervation as they were not expressed by body myonuclei and their expression were maintained at similar levels in NMJ myonuclei. We suggest that continued agrin-Lrp4/MuSK signaling might be the source for maintaining their expression. To test this hypothesis, we deleted *Musk* using the AAV-CRISPR/Cas9 system. Three weeks post-sgRNA injection, when *Musk* transcript was significantly downregulated, ∼18% of these “denervation resistant” synaptic genes were downregulated (Fig. 3e). Some of the most strongly reduced transcripts were *Prkar1a*, *Ryr3* and *Colq*, supporting the idea that MuSK is an important regulator of synaptic gene expression ^62, 63^. However, ∼82% of the “denervation resistant” synaptic genes remained unchanged after *Musk* knockout. While we cannot exclude that three weeks were not sufficient to completely abrogate agrin-Lrp4/MuSK signaling, our results also suggest that other neurotrophic factors, including some that have previously been implicated in the regulation of synaptic gene expression (e.g., neuregulin-ErbB and Wnt signaling), may maintain synaptic gene expression ^64, 65, 66, 67, 68^.

To get further insights into the molecular mechanisms of NMJ maintenance, we tackled the function of *Etv4, Lrtm1* and *Pdzrn4*, three not-described NMJ-specific genes. We used AAVMYO-mediated overexpression and a muscle fiber-specific AAVMYO-CRISPR/Cas9 knockout system, which has been described to copy the phenotype of gene knockout mice ^40^. ETV4 overexpression was sufficient to trigger a synapse-specific gene transcription signature in body myonuclei by upregulating more than half of the NMJ-myonuclei transcripts (Fig. 5). These results indicate that a large portion of the synaptic genes are regulated through the ‘N-box’ domain ^69^. However, depletion of ETV4 in adult skeletal muscle did not affect NMJ structure. A likely reason could be compensation by other ETS transcription factors including ETV5 and GABP, which are both expressed in skeletal muscle. Interestingly, whole-body knockout of *Etv5* results in an NMJ phenotype ^70^. *Etv5* is expressed at much higher levels than *Etv4* (based on LCM data ^31^) and as both can bind the same ‘N-box’ domain ^71^, *Etv4* may be redundant for NMJ maintenance in the presence of *Etv5*. Similarly, knockout of *Lrtm1* did not significantly affect muscle function and NMJ morphology and overexpression of LRTM1 also did not have a strong effect on the gene expression pattern in muscle. We therefore conclude that LRTM1 may also be compensated for by similar proteins, as there are many other transmembrane proteins with extracellular leucine-rich repeats ^44^. Another possibility is that LRTM1 does not play a role in NMJ maintenance as we hypothesized but rather acts as an auxiliary protein for potassium channels. A group of leucine-rich repeat-containing membrane proteins have been observed to cause negative shifts in potassium channel voltage dependence ^72^. It is also important to note that the AAV-CRISPR/Cas9-mediated knockout system tests for the necessity of a gene in NMJ maintenance and not for its function during NMJ development. Whole-body knockouts might reveal more drastic phenotypes as they interfere with gene function during development.

PDZRN4 overexpression caused partial mislocalization of AChRs and MuSK (Fig. 5c, Supplementary Fig. S5c) and knockout induced NMJ fragmentation (Fig. 6g). Based on these observations and the ability of PDZRN3, a paralogous protein of PDZRN4, to bind MuSK ^45^, we hypothesized that PDZRN4 may also be a binding partner of MuSK. Indeed, PDZRN4 co-immunoprecipitated with MuSK when expressed in HEK 293T cells (Fig. 7). Interestingly, removal of the PDZ domain of PDZRN4 did not abolish MuSK interaction, setting it apart from PDZRN3. Based on PDZRN4 localization at the Golgi apparatus and the accumulation of MuSK at the Golgi in HeLa cells when overexpressed together with PDZRN4, we propose a model where PDZRN4 acts as a tether between the Golgi and MuSK. The Golgi apparatus acts as a protein trafficking center, where proteins are either sent to the plasma membrane or sorted for degradation ^73^. Additionally, the Golgi apparatus is a site of post-translational modifications as proteins can be acetylated, sulfated, phosphorylated, palmitoylated, O-glycosylated and N-glycosylation sites can be modified ^74^. PDZRN4 may prolong the time MuSK spends at the Golgi apparatus allowing post-translational modifications of its N-glycosylation sites ^75^. Notably, overexpression of PDZRN4 resulted in the appearance of an additional band for MuSK that migrated at a lower apparent molecular mass in Western blots. Interestingly this low molecular mass version of MuSK preferentially bound to PDZRN4 (Fig. 7e, Supplementary Fig. S7e). However, whether improper post-translational modification of MuSK is sufficient to cause NMJ fragmentation in *Pdzrn4* knockout mice remains elusive, as changes in MuSK glycosylation did not affect MuSK function in cultured myotubes ^75^. It is possible that PDZRN4 is also involved in the post-translational modification of proteins other than MuSK, as AChRs appeared in the same mislocalized MuSK vesicles (Fig. 5c, Supplementary Fig. S5c). Another indicator that PDZRN4 might tether other synaptic proteins at the Golgi apparatus is that the same *Pdzrn4* splice version is also expressed in different neurons in the brain ^76^. Importantly, the NMJ-specific expression of *Pdzrn4* can also be observed in human skeletal muscle snRNA-seq data ^77^.

In summary, we perform snRNA-seq on skeletal muscle and identify numerous, previously undescribed transcripts in NMJ myonuclei. We studied synaptic gene expression in NMJ and body myonuclei in denervation and find that synaptic gene expression relies not only on neurotrophic factors but also on electrical activity. Collectively, our snRNA-seq data provide a valuable resource which can be explored online on our interactive atlas (https://ruegglab.shinyapps.io/snrnaseq/). Furthermore, we characterize *Etv4, Lrtm1* and *Pdzrn4* in muscle by overexpression and knockout and identified PDZRN4 as an interactor of MuSK protein at the Golgi apparatus, suggesting a novel pathway for NMJ post-translational modifications.

## Methods

### Mice

The mice were kept at the local animal facility (Biozentrum, Basel) under a 12-hour light to dark cycle, with *ad libitum* access to standard laboratory chow and water. All animal studies were performed in accordance with the law and guidelines of the Swiss authorities and were approved by the veterinary commission of canton Basel-Stadt (Schlachthofstrasse 55, 4056 Basel, Switzerland). All mice used in the experiments were 2-14 months of age. Within an experimental group, the maximum age difference between mice was ± 40%. Muscle-specific Cas9 mice (Cas9mKI) were generated by crossing Cas9 knock-in mice ^78^ with HSA-Cre mice ^79^.

### Muscle function

Grip strength was measured using a force meter (Columbus Instruments) attached to either a triangular bar (for forelimbs) or a grid (for all limbs). Mice were held close to the bar or on top of the grid until they grasped it firmly and were then gently pulled away horizontally at a constant speed until they lost their grip. The measurement was repeated at least five times with short breaks in between, and the median of all values was used as the final performance.

For the balance beam experiments, the mice were trained one day prior to the actual tests. The mice were placed on a small platform where they learned to walk up across a 12 mm wide square metal bar to another platform with a red Plexiglas house. The next day, the mice were again placed on the small platform and the time to cross a 10 mm bar over a set 80 cm distance was measured. This was repeated three times with 30 s rest periods in between. The best value over the three trials for each mouse was used as the final performance.

### CMAP measurements

CMAP was performed similar to a previous publication ^31^. Briefly, mice were anaesthetized and the sciatic nerve was stimulated by trains of 10 stimulations of increasing frequencies ranging from 3 Hz to 100 Hz. Action potentials were recorded using a needle electrode inserted into the GAS muscle, with a reference electrode placed subcutaneously for grounding. Potential decrements after repetitive stimulations were quantified by measuring the change between the first stimulation and subsequent ones.

### Surgical denervation and BoTX injection

Mice were anaesthetized by isoflurane inhalation and a small incision on the skin near the sciatic nerve was made. The sciatic nerve was raised using a glass hook and a small 5 mm piece was excised. Surgical clips were used to close the wound and the mice were returned to their cage. The mice received analgesia (Buprenorphine, 0.1 mg/kg) 1 h before, 4-6 h after and twice on the next day of the surgery.

When denervated by botulinum toxin (BoTX), mice were anaesthetized by isoflurane inhalation and received an intra-muscular injection of 25 µl Dysport® (40 U/ml in saline solution) into the TA, as previously described ^80^. BoTX control mice were injected with 25 µl of saline solution.

### AAV administration

For intra-muscular injections, one leg was shaved with an animal trimmer and the region of the TA was disinfected. For overexpression experiments, AAVs were diluted in PBS to a concentration of 5 x 10^11^ vector genome copies (vg) per ml and 50 µl (i.e. 2.5 x 10^10^ vg) were injected into the TA muscle. To knockout *Musk* in mouse muscle for snRNA-seq, 50 µl of 1.5 x 10^12^ vg/ml (i.e. 7.5 x 10^10^ vg) were injected into the TA. To generate systemic knockouts, mice were injected with 100 µl of 3 x 10^12^ vg/ml (i.e. 3 x 10^11^ vg) via the tail vein. All injections were made with a 30 G 0.3 ml insulin syringe and the mice were anaesthetized by isoflurane inhalation.

### Single guide RNA design and vector assembly

The sgRNAs targeting *Musk* were designed and tested previously ^40^. The single guides targeting *Etv4*, *Lrtm1* and *Pdzrn4* were selected using CRISPOR ^81^ and Geneious Prime® (version 2020.2.3). Six to seven different guides were assembled into one plasmid using a multiplex CRISPR/Cas9 assembly method ^82^. The guides were then cloned into an AAV transfer vector containing ITRs, as described previously ^40^.

### AAV production and purification

AAVs were produced as previously described ^40^. Briefly, HEK293T (ATCC CRL-3216) cells were transfected with AAVMYO ^41^, pAdDeltaF6 helper (a gift from J. M. Wilson, Addgene, #112867) and an AAV transfer plasmid containing an overexpression construct or sgRNA expression cassettes. Two days’ post-transfection the cell culture medium containing AAVs was collected. Three days’ post-transfection HEK293T cells were centrifuged and the supernatant was pooled with the cell culture medium from the previous day. HEK293T cells were lysed with AAV lysis solution (50 mM Tris-HCl, 1 M NaCl, 10 mM NgCl2, pH 8.5) and 50 U of salt active nuclease (SRE0015-5KU, Sigma) for 1 h at 37°C. The lysate was centrifuged at 3200 g, 15 min at 4 °C and the supernatant containing AAVs was collected. The cell culture medium containing AAVs was mixed with polyethylene glycol 8000 (89510, sigma) to a concentration of 8% (w/v) and incubated for 2 h at 4 °C followed by centrifugation at 4000 g, 30 min at 4 °C. The AAV pellet was resuspended with 1 ml AAV lysis solution and pooled with the cell lysate. To purify AAVs the entire lysate was loaded onto a 15-25-40-60% iodixanol (Serumwerk) gradient and centrifuged at 65’000 rpm (Beckman type 70 Ti rotor) for 2 h at 4 °C. Purified AAVs were collected from the 40% iodixanol phase and extracted via buffer exchange using a 100 kDa MWCO filter (Millipore). The concentration of the virus was measured by qPCR using forward (fw) primer GGAACCCCTAGTGATGGAGTT and reverse (rv) primer CGGCCTCAGTGAGCGA targeting AAV2 ITRs as previously described ^83^.

### Single nuclei isolation for snRNA-seq

Every step of single nuclei isolation was performed on ice or at 4 °C. The nuclei were always isolated from freshly dissected muscle that was kept in ice cold PBS for maximum 1 h after dissection. GAS snRNA-seq results were obtained from one muscle and TA results were always a pool of two TA muscles. The tissue was individually minced with scissors (14084-08, FST) for 3 min in 2 ml microcentrifuge tubes together with 200 µl single nuclei lysis buffer (250 mM sucrose, 25 mM KCl, 4 mM MgCl_2_, 10 mM Tris-HCl pH 8 and 0.1% NP40 in nuclease free water). The minced muscle was transferred to a 7 ml dounce homogenizer (D9063, Sigma-Aldrich) with additional 3 ml single nuclei lysis buffer and carefully dounced with 8-10 strokes. The muscle slurry was transferred to a 50 ml falcon tube and the dounce homogenizer was rinsed with 3 ml single nuclei wash buffer (1% BSA in PBS) which was pooled with the previous fraction. The muscle slurry was filtered through 70 µm (352350, Falcon) and 30 µm (130-098-458, Miltenyi Biotec) cell strainers to remove large pieces of tissue and cell debris. The filtered lysates were then centrifuged for 5 min at 500 g in 50 ml falcon tubes and the supernatant was removed leaving ∼200-300 µl liquid in the tube. The pellet was resuspended with 1 ml of single nuclei wash buffer containing 0.2 U/µl RNase inhibitor (3335402001, Roche) and transferred to a microcentrifuge tube. After adding 1 µl Hoechst solution the single nuclei suspension was centrifuged again for 5 min at 500 g. Next, the entire supernatant was removed and the nuclei pellet was resuspended in 100-400 µl of single nuclei wash buffer with RNase inhibitor depending on the amount of starting tissue. The quality and concentration of nuclei was controlled on a fluorescent microscope using a hemocytometer (concentration would optimally range from 500-1500 nuclei/µl).

### Single nuclei library construction and sequencing

To generate single nuclei libraries 10x Genomics Chromium Next GEM Single Cell 3’ kits (PN-1000121, PN-1000120, PN-1000213) and the corresponding protocol were used. We always aimed for 10’000 nuclei libraries per sample which is the maximum recommended by the manufacturer. For the most part the manufacturer’s instructions were followed, the only variables that had to be adjusted were the cDNA amplification and sample index PCRs (steps 2.2 and 3.5 respectively) where 12 cycles were used in both cases. Quality control was performed on an Agilent Fragment Analyzer using the HS NGS Fragment Kit (1-6000 bp). cDNA ranging from 300 to 3000 bp with a peak around 1000 bp was considered good quality. After library construction samples were quality controlled again and were sequenced paired end (PE) 28/91 on a NovaSeq 6000 system (Illumina). We aimed to reach >30’000 reads per nucleus for all samples. Sequencing and quality control was performed by the genomics facility at the D-BSSE (ETH, Basel, Switzerland).

### Single nuclei RNA-seq alignment, clustering and analysis

Demultiplexing was performed with bcl2fastq (version 2.20) and was done by the genomics facility at the D-BSSE (ETH, Basel, Switzerland). FASTQ files were aligned to mouse reference mm10 (GENCODE vM23/Ensembl 98, from 2020) using cellranger count version 5.0 with the argument – include-introns. As a quality control measure we looked at the percentage of reads mapped to intronic regions which should range around 50-60% for nuclei.

Cellrangers output folders “filtered_feature_bc_matrix” were loaded into R (version 4.0.3) using Seurats (version 4.0.0) Read10x function. Nuclei expressing fewer than 500 features and more than 5% of mitochondrial features were removed. Next, nuclei were clustered following the fast integration (RPCA) workflow (https://satijalab.org/seurat/articles/integration_rpca). Any low quality clusters (low feature count and high expression of mitochondrial genes) or multiplets (co-expression of several cell type specific markers) were removed manually. After removal, the remaining nuclei were normalized and clustered again using RPCA with SCTransform normalization. For differential expression between clusters and between conditions first NormalizeData() was used on the “RNA” assay and then the FindMarkers() function was run. For visualization Seurats DimPlot(), FeaturePlot(), VlnPlot() and DotPlot() were used.

To generate our list of synaptic transcripts we compared the NMJ cluster to clusters Type IIA, Type IIX and Type IIB. To be more stringent we set a threshold for the adjusted p-value to < 5E-5 and the log2FC to ±0.25. The same procedure was done to generate a list of differentially expressed genes for the MTJ cluster. To see which transcripts are changed in denervation (by nerve cut), after BoTX injection and *Musk* knockout, the FindMarkers() function was used and we set a threshold for the p-value to < 0.01 and the log2FC to ±0.25.

### RNA isolation and bulk-RNA-seq analysis

TA muscles were crushed to powder on a metal block cooled with liquid nitrogen. Total RNA was extracted from the muscle powder using the RNeasy Fibrous Tissue Mini Kit (74704, Qiagen) according to the manufactures protocol. RNA concentration and integrity were measured using a Quant-iT^TM^ RiboGreen RNA assay kit (Invitrogen) and a Bioanalyzer (Agilent), respectively. Libraries were prepared using the TruSeq stranded mRNA library kit (20020595, Illumina) starting from 200 ng of RNA. Sequencing was performed on the NovaSeq 6000 (Illumina) system (PE 2×51) which resulted in 26-52 mio reads per sample. FASTQ files were aligned to the indexed mouse transcriptome mm10 using Salmon (version 1.1.0) with the flags validateMappings, seqBias and gcBias. The output quant.sf files from all samples were imported into R (version 4.1.2) using tximeta (Bioconductor) and analyzed with DESeq2 (Bioconductor). Transcript level information was summarized to gene level and genes with fewer than 5 counts across all samples were removed. The design was set to “∼ Injection” and after applying DeSeq() the result for each group was exported individually together with PBS-injected controls. All genes with adjusted p-values < 0.05 and Log2FC ± 0.25 were considered as significantly changed.

### Sashimi plots and bulk RNA-seq exon coverage

To generate sashimi plots bulk RNA-seq FASTQ files from TA and *soleus* muscles from 10 month old male mice were downloaded from GEO (GSE139204) [9]. The FASTQ files were aligned to mouse reference GRCm39/mm39 using STAR (version 2.7.9a) and the genomic region of each gene was extracted from bam files using samtools (version 1.12). The extracted region was imported into Integrative Genomics Viewer (IGV, version 2.16.1) and Sashimi plots were generated. The Sashimi plots were extracted and adjusted for better visualization.

### qPCR

Total RNA was isolated as described above. 500 ng of RNA were used to generate cDNA with the iScript cDNA Synthesis Kit (Bio-Rad). The cDNA was diluted 1:5, mixed with PowerUp SYBR Green Master Mix (Thermo Fisher) and combined with the corresponding primers at a final primer concentration of 500 nM. The qPCR reactions were run on a QuantStudio^TM^ 5 PCR system (Thermo Fisher). For quantification all targets were normalized to housekeeper *Gapdh*. The primer sequences used were: *Gapdh* fw: ACC CAG AAG ACT GTG GAT GG, rv: GGA TGC AGG GAT GAT GTT CT; *Etv4* fw: AGG AGT ACC ATG ACC CCC TG, rv: GGA CAT CTG AGT CGT AGG CG; *Lrtm1* fw: CCC TTG GAT TTG TGA CTG CC, rv: GAG GAA TGG AGA GGA GAG CC; *Pdzrn4* fw: CTT AGA CTC CGA CTG TGC CA; rv: GCA CTC GAT GGT ACT GCT TG; *Musk* fw: GCT CCT GAA TCC CAC AAT GTC, rv: AGA GTC CTG GCT TTG TGA TGA.

### Immunofluorescence staining on cross-sections

Muscles were embedded in OCT, frozen in isopentane and cut to 10 µm sections using a cryostat. Sections were fixed with 4% PFA in PBS for 5 min followed by neutralization with 100 mM glycine in PBS for 2 x 10 min. Next, sections were permeabilized and blocked with blocking solution (3% BSA, 0.4% Triton X-100 in PBS) for 30 min. Primary antibodies were diluted in 3% BSA in PBS and added to the sections for either 1 h at room temperature or overnight at 4 °C (for antibodies and dilutions see Supplementary Table 1). This was followed by three washing steps with PBS. Secondary antibodies and DAPI were diluted in 3% BSA in PBS and added to the samples for 1 h in the dark. To finish, samples were washed with PBS and mounted with ProLong^TM^ Gold antifade (Invitrogen). Images were taken on either a wide field IX83 microscope (Olympus) or a confocal LSM 700 (Zeiss).

### HeLa cell transfection and staining

HeLa cells (ATCC CCL-2) were cultured in DMEM (D5796, Sigma) containing 10% FBS, 1% Penicillin/Streptomycin and 0.01 M HEPES. For IF imaging HeLa cells were cultured on 4-well cell culture chamber slides with removable frames (Sarstedt). Transfection was performed using Lipofectamine 2000 (Invitrogen) where all wells received 260 ng/cm^2^ of DNA and 1.04 µl/cm^2^ of Lipofectamine. 24 h after transfection HeLa cells were washed with PBS and fixed for 15 min with 4% PFA in PBS. IF stainings were performed as described above. For antibodies and dilutions see Supplementary Table 1.

### Whole mount immunostaining

EDL muscles were carefully dissected leaving the tendons on both ends intact. Muscles were pinned on a sylgard dish and fixed with 4% PFA in PBS for 30 min at room temperature, followed by three washes with PBS for 5 min each. Muscles were stored in 50% glycerol in PBS (v/v) at −20 °C until usage. Fixed muscles were cut into small bundles and permeabilized with 2% Triton X-100 in PBS for 2 h. Muscle bundles were then neutralized with 100 mM glycine in PBS and blocked in blocking solution (3% BSA, 0.2% Triton X-100 in PBS) for 30 min each. This was followed by primary antibody incubation in blocking solution overnight at 4 °C (for antibodies and dilutions see Supplementary Table 1). On the next day samples were washed with PBS containing 0.2% Triton X-100 four times and incubated in secondary antibody solution and α-bungarotoxin for 2 h. Samples were washed again four times in PBS containing 0.2% Triton X-100 and then mounted with ProLong^TM^ Gold antifade (Invitrogen). Images were taken at the Imaging Core Facility (Biozentrum, Basel, Switzerland) using a confocal LSM 700 (Zeiss) at 63x magnification. 10-15 individual NMJs were imaged and quantified for each mouse and the average value was used for statistics.

### Single molecule fluorescent in situ hybridization (smFISH)

For detection of RNA on cross-sections we used the V1 kit from RNAscope (ACDBio) and followed the manufacturers protocol. Briefly, fresh frozen 10 µm muscle cross-sections were fixed with 4% PFA for 15 min on ice, followed by an ethanol series of 50, 70 and 100% at room temperature to dehydrate the tissue. Next, muscle sections were digested with protease IV for 30 min, or 5 min if combined with an IF staining. After washing with PBS the cross-sections were placed in a humid chamber with hybridization probes followed by several amplification steps. The sections were stained with DAPI and mounted with ProLong^TM^ Gold antifade (Invitrogen). Images were taken at the Imaging Core Facility (Biozentrum, Basel) using a confocal LSM 700 (Zeiss).

### Differential fractionation and western blot

TA muscles were powdered on a cooled metal block and lysed in 400 µl cytosol lysis buffer (50 mM Tris-HCl pH 7.5, 150 mM NaCl) supplemented with protease inhibitors (11 836 170 001, Roche) for 30 min at 4 °C. Lysates were centrifuged at 100’000 g for 30 min at 4 °C in a micro ultracentrifuge (CS 150FNX, Hitachi). The supernatant (cytosolic fraction) was transferred to a new tube and the pellet was resuspended with 300 µl membrane lysis buffer (50 mM Tris-HCl pH 7.5, 150 mM NaCl, 1% NP-40) supplemented with protease inhibitors (Roche). After 20 min of incubation the samples were centrifuged again at 100’000 for 30 min at 4 °C. The supernatant (membrane fraction) was transferred to a new tube. Protein concentrations of both the cytosolic and membrane fractions were measured using the Pierce™ BCA Protein Assay Kit (Thermo Fisher Scientific) and equal amounts of protein were loaded on to a NuPAGE 4-12% Bis-Tris gel (Invitrogen). Western blots were prepared and imaged as previously described ^84^.

### Immunoprecipitation

HEK 293T cells were cultured in 6-well plates in DMEM (D5796, Sigma) containing 10% FBS, 1% Penicillin/Streptomycin and 0.01 M HEPES. Transfection was performed using Lipofectamine 2000 (Invitrogen) where all wells received 260 ng/cm^2^ of DNA and 1.04 µl/cm^2^ of Lipofectamine. The cells were transfected with *Musk*, *Pdzrn4-Flag* and *pac* (control) plasmids, all containing a CMV promoter. 24 h after transfection cells were washed in PBS and lysed with 500 µl IP lysis buffer (20 mM Tris-HCl pH 7.4, 100 mM NaCl, 5 mM EDTA, 0.5% NP-40) supplemented with protease (Roche) and phosphatase inhibitors (Roche). The lysed cells were transferred to microcentrifuge tubes, sonicated for 3 s and incubated on a rotating wheel for 30 min at 4 °C. The lysates were filtered through 70 µm cell strainers before measuring protein concentration using the Pierce™ BCA Protein Assay Kit. 500 µg of protein was diluted to a concentration of 1 µg/µl and 2 µg of anti-FLAG M2 antibody (Sigma) was added and incubated overnight at 4°C for IP. Antibodies were pulled down using magnetic Dynabeads^TM^ Protein G (10004D, Invitrogen) and washed with IP lysis buffer. The pulled down proteins were released from the magnetic beads by adding 50 µl 1x Laemmli buffer and heating to 95°C for 10 min. Equal volumes were loaded onto NuPAGE 4-12% Bis-Tris gels (Invitrogen) together with whole protein lysates. Western blots were prepared and imaged as previously described ^84^.

## Acknowledgements

We thank the Imaging Core Facility, sciCORE, and Animal facility of the Biozentrum for their support with imaging, computing and mouse housing, respectively. We acknowledge Elodie Burcklen, Phillippe Demougin and Christian Beisel of the Genomics facility of the D-BSSE (ETH) for their support in library preparation, quality control and sequencing. We thank D. J. Ham for his advice on the manuscript and for helping acquire funding. This work was funded by the Swiss National Science Foundation (#189248) and the cantons of Basel-Stadt and Basel-Landschaft awarded to M.A.R.

## Author Contributions

A.S.H. and M.A.R. conceptualized the study; A.S.H. developed the methodology; A.S.H., S.L., A.T., M.T., and F.O. performed the experiments, A.S.H. and S.L. quantified the data; A.S.H. analyzed the data, A.S.H. and M.A.R. interpreted the data and wrote the manuscript; M.A.R. acquired the funding.

---

**Figure S1:**
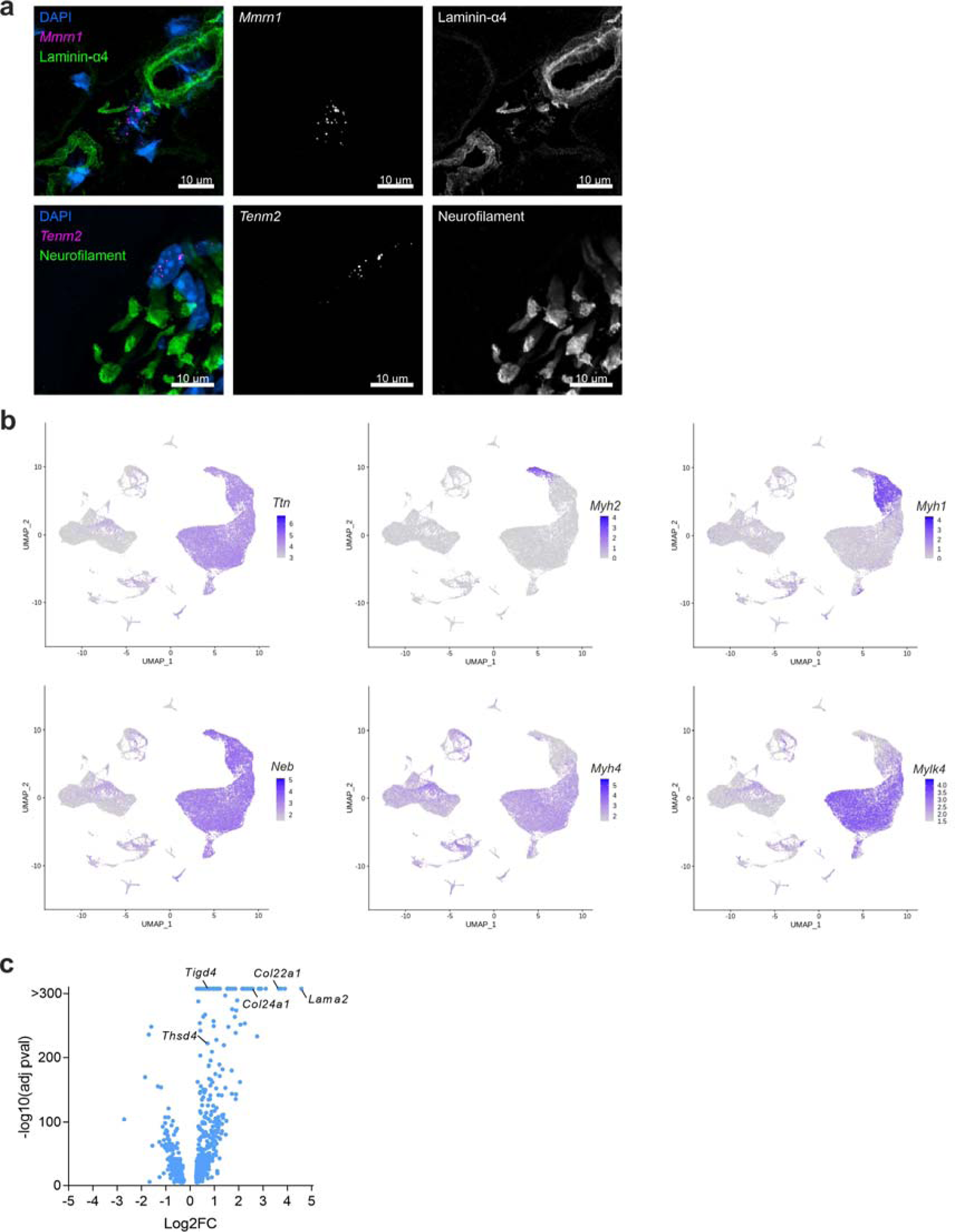
Additional data supporting the control skeletal muscle UMAP analysis. **a** Single molecule fluorescent *in situ* hybridization (smFISH) of *Mmrn1* or *Tenm2* combined with an immunofluorescence staining of Laminin α4 or neurofilament on *gastrocnemius* (GAS) muscle. *Mmrn1*-expressing cells are located in close proximity to larger blood vessels and *Tenm2*-expressing cells are located near axons. **b** Feature plots visualizing muscle fiber specific and muscle fiber type specific markers. **c** Volcano plot of differentially expressed transcripts in MTJ myonuclei compared to body myonuclei (max. padj. cutoff at 2.23E-308).

**Figure S2:**
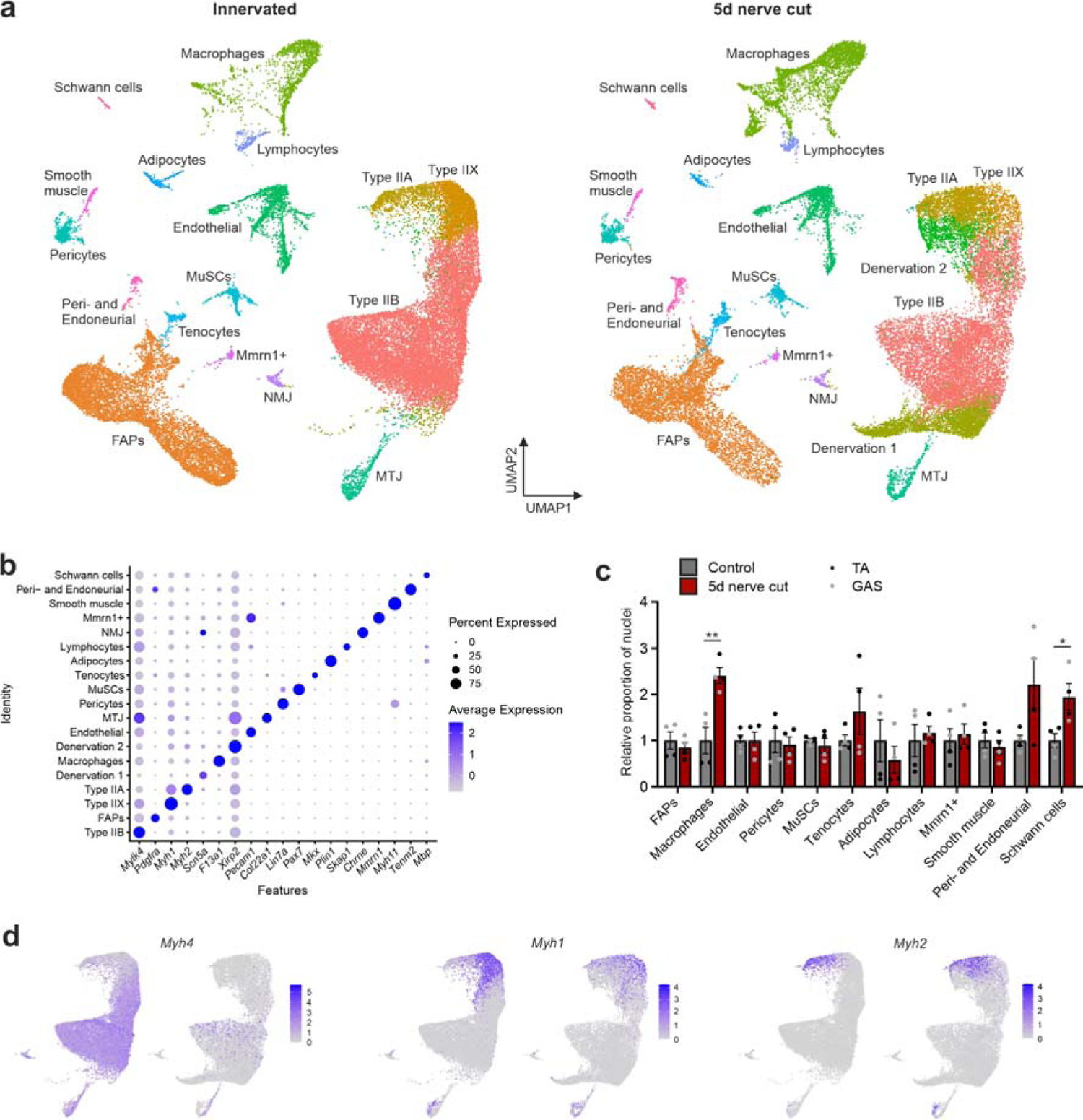
Sciatic nerve cut UMAP and supporting data. **a** Split UMAP of ∼65’000 nuclei derived from five day denervated and control muscles. *n* = 2 for TA and Gast for both conditions. **b** Dot-plot visualizing the genes used to assign cell types to clusters in the UMAP in (**a**). **c** Relative proportion of nuclei of all mononucleated cell types in TA and GAS and the effect of denervation. **d** Feature plots of myonuclei visualizing the expression of different myosin heavy chain isoforms in control versus denervated muscle. In (**c**) the data is presented as mean ± SEM and a regular t-test was performed (p < 0.05 = *, p < 0.01 = **, p < 0.001 = ***).

**Figure S3:**
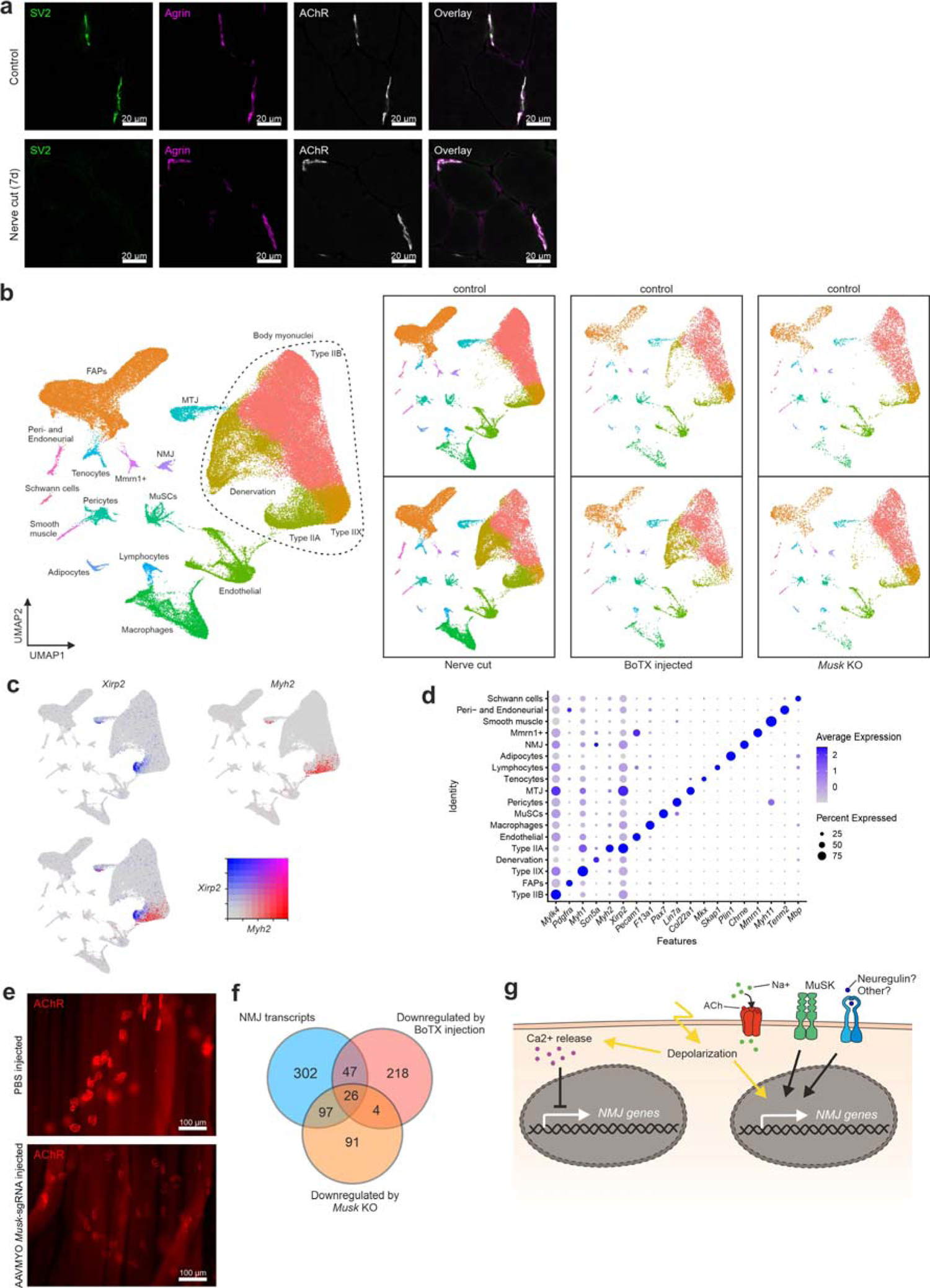
Nerve cut, BoTX injected and *Musk* KO supporting data. **a** IF staining of NMJs on TA cross-sections 7 days after sciatic nerve transection and in control muscle. AChRs were visualized using α-bungarotoxin (BTX) and nerve terminals by IF staining against synaptic vesicle protein 2 (SV2). Note that agrin remains bound at the synaptic basal lamina and that AChRs remains clustered at the NMJ despite loss of motor neuron axons (lack of SV2). **b** UMAP depicting ∼100’000 nuclei isolated from TA and GAS muscles after nerve cut, BoTX injection and MuSK depletion (*Musk* KO). The different conditions and respective controls are additionally visualized as split UMAPs to the right. **c** Dual gene feature plot depicting a *Myh2* devoid region within the type IIA cluster expressing the denervation 2 marker *Xirp2* (see Fig. 2). **d** Dot-plot visualizing the genes used to assign cell types to clusters in the UMAP in (**b**). **e** Whole mount preparation of TA muscles injected with either PBS or AAVMYO encoding *Musk*-sgRNA. Reduced BTX staining (AChR) confirms successful reduction of MuSK protein. **f** Venn diagram visualizing how many NMJ transcripts are downregulated in NMJ myonuclei after BoTX injection or *Musk* KO. **g** Illustration of which processes promote synaptic gene expression in NMJ myonuclei. In (**f**) transcripts were categorized as significantly changed when p-val. < 0.01 and log2FC ±0.25.

**Figure S4:**
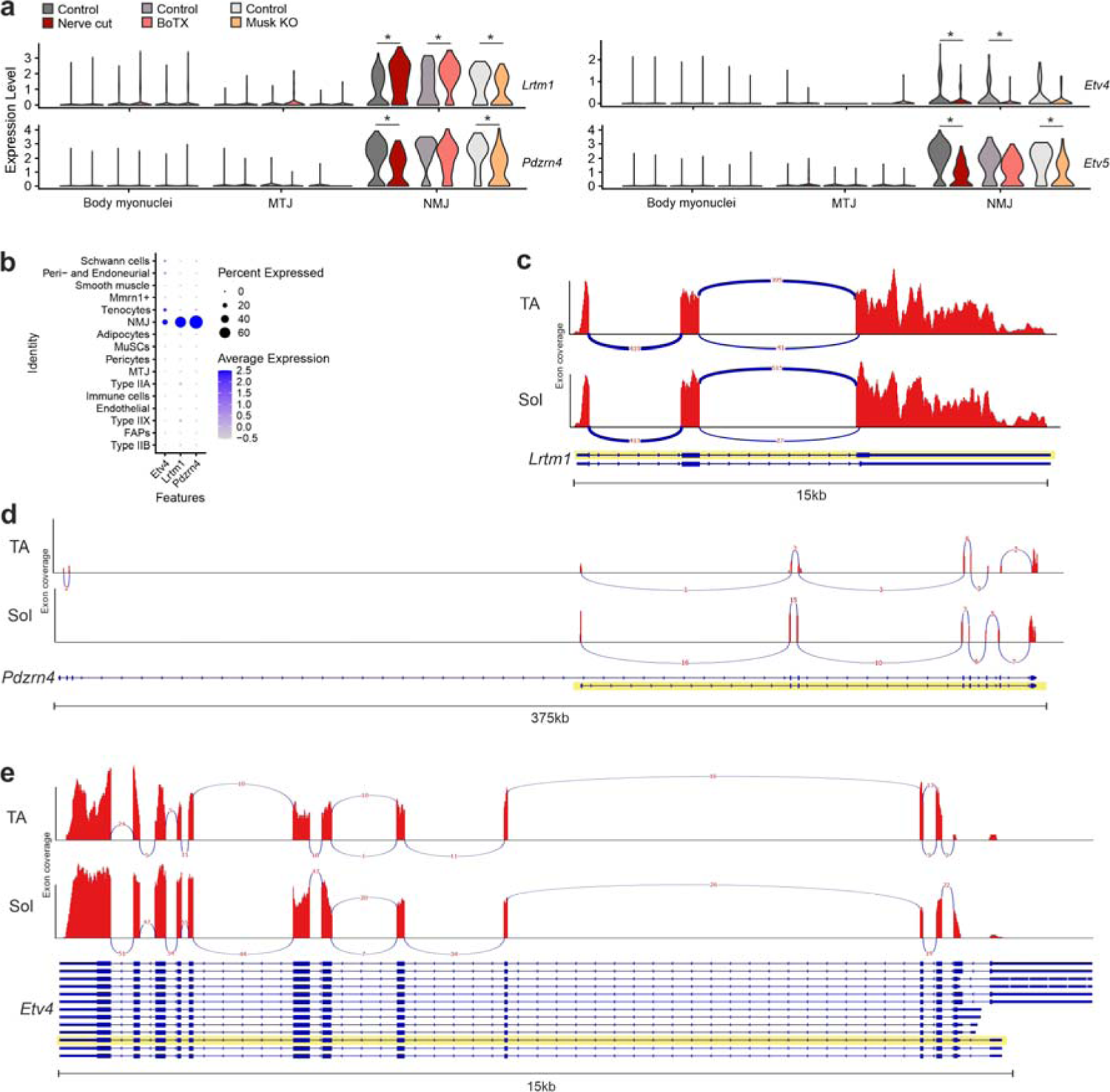
Data supporting NMJ gene target selection. **a** Violin plots generated from the snRNA-seq data shown in Fig. 3a. **b** Dot-plot generated from the snRNA-seq data depicted in Fig. 1a showing expression of *Etv4*, *Lrtm1* and *Pdzrn4* in the different nuclei populations in innervated muscle. **c**, **d** and **e** bulk RNA-seq reads from TA and *soleus* (Sol) for *Lrtm1* (**c**), *Pdzrn4* (**d**) and *Etv4* (**e**) were extracted and visualized as Sashimi plots using integrative genomics viewer. The numbers between splice junctions show the number of reads mapped between splice-sites. The dominant splice isoform is highlighted in yellow. The NCBI nucleotide ID is NM_176920 for *Lrtm1*, NM_001164594 for *Pdzrn4* and NM_001316365 for *Etv4*.

**Figure S5:**
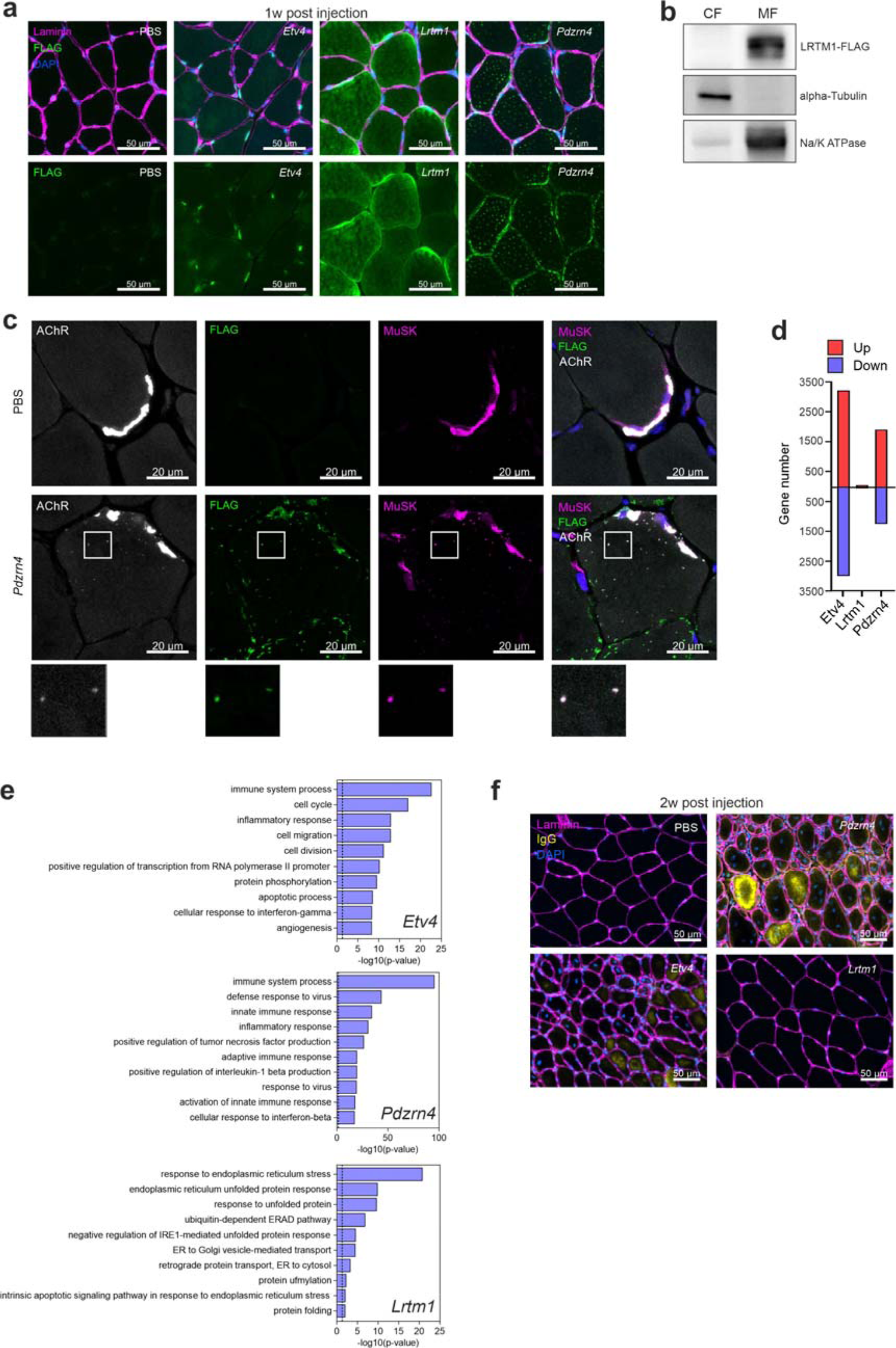
Overexpression of *Etv4*, *Lrtm1* and *Pdzrn4* in mouse muscle. **a** Immunofluorescence staining against the FLAG-tag on TA cross-sections one-week post AAVMYO or PBS injection. ETV4 localizes to myonuclei, LRTM1 is mainly found at the muscle fiber periphery and PDZRN4 is detected near the sarcolemma and in vesicle-like structures within muscle fibers. **b** Western blot analysis of a TA muscle overexpressing LRTM1 that was separated into a cytosolic fraction (CF) and a membrane fraction (MF). Alpha tubulin and Na/K ATPase are positive controls. **c** Immunofluorescence staining on TA cross-sections showing co-localization of AChRs and MuSK within the vesicle-like structures positive for PDZRN4. **d** Number of differentially expressed genes in bulk RNA-seq one-week post AAVMYO injection (padj < 0.05, Log2FC ± 0.25). **e** Top 10 DAVID GO-terms (biological processes) associated with upregulated genes in the indicated samples. **f** Immunofluorescence staining against mouse IgG on TA cross-sections two weeks after AAVMYO or PBS injection. ETV4 and PDZRN4 overexpression causes de- and regeneration of muscle fibers, indicated by the presence of centralized nuclei and of IgG-positivity of muscle fibers.

**Figure S6:**
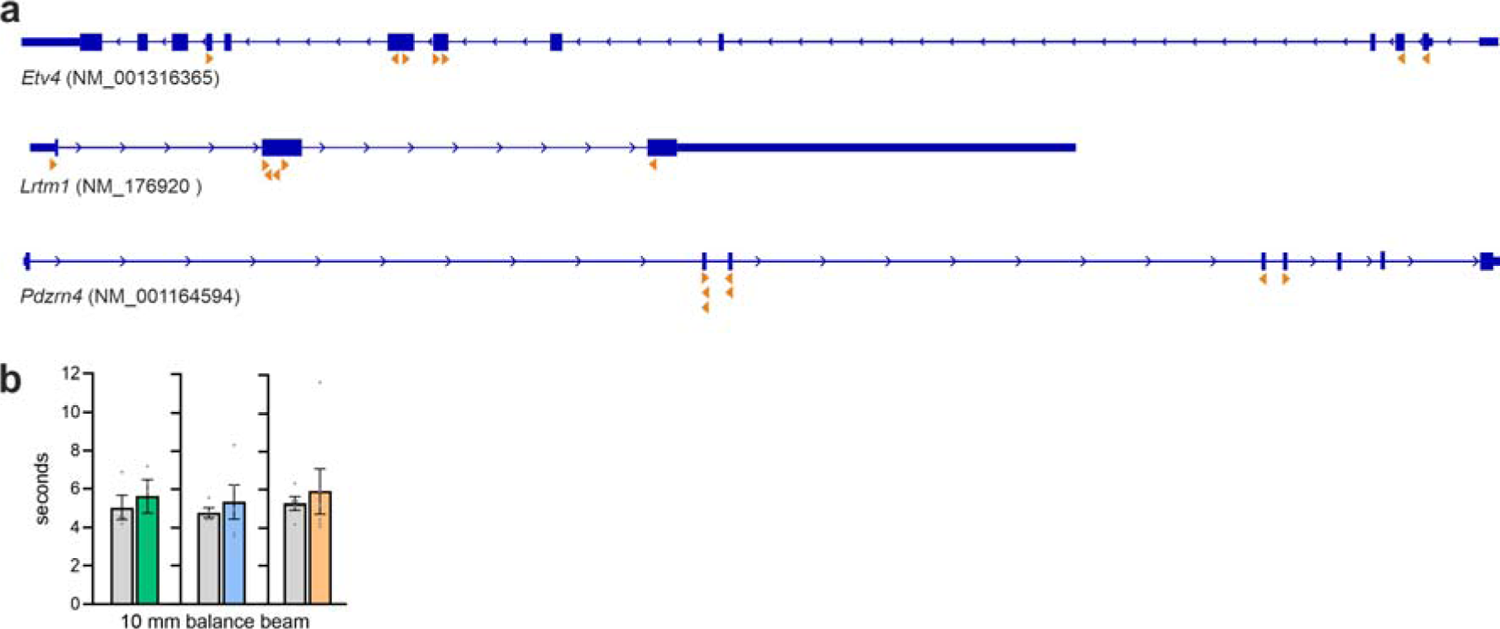
Single guide RNA localization and balance beam experiment. **a** Exon map with orange arrow heads indicating the location and direction of each single guide RNA. **b** Balance beam performance of knockout mice. After an initial training to walk across a 12 mm narrow metal bar, mice were timed to traverse a 10 mm balance beam, covering a distance of 80 cm.

**Figure S7:**
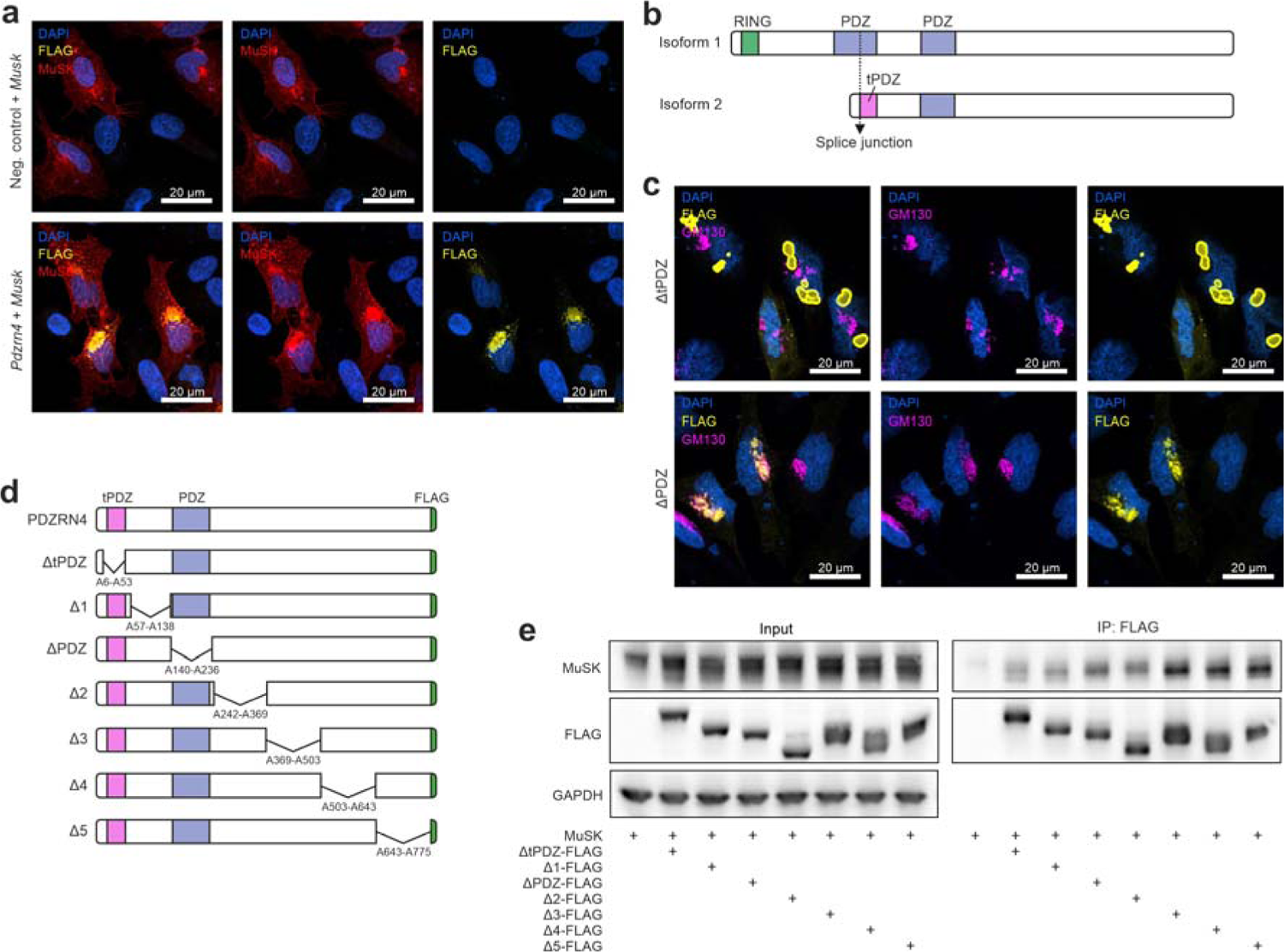
PDZRN4 localizes to the Golgi complex in HeLa cells and binds to MuSK. **a** Overexpression of PDZRN4 and MuSK in HeLa cells. HeLa cells transfected with expression constructs for MuSK (top) or co-transfected with PDZRN4-FLAG-encoding plasmids (bottom) were stained for FLAG, MuSK and DAPI. Co-expression of PDZRN4 enhances MuSK immunoreactivity in the Golgi. **b** Schematic of the two PDZRN4 isoforms. Skeletal muscle only expresses isoform 2 that lacks the RING domain and contains a truncated PDZ domain (termed tPDZ). **c** IF staining on HeLa cells overexpressing different deletion mutants of PDZRN4. The mutant lacking tPDZ does not localize to the Golgi while the ΔPDZ mutant show a Golgi localization like full-length PDZRN4. **d** Schematic of deletion mutants of PDZRN4. **e** Co-immunoprecipitation experiments using deletion mutants expressed in HEK 293T cells together with MuSK using antibodies against the FLAG-tag. MuSK is present in all precipitates to a different degree.

**Supplementary Table 1:** List of all antibodies and dilutions used in experiments.

| Figure | Method | Antibody target | Host | Source | Dilution |
| --- | --- | --- | --- | --- | --- |
| Fig. S1a | RNAscope + Immunostaining | Laminin α4 | Chicken | Gift from P. Yurchenco | 1:400 |
| Fig. S1a | RNAscope + Immunostaining | Neurofilament | Rabbit | Sigma (N4142) | 1:5'000 |
| Fig. S3a | Immunostaining | Synaptic vesicle protein 2 | Mouse | DSHB | 1:300 |
| Fig. S3a | Immunostaining | Agrin | Rabbit | Self made (C-Term 204) | 1:2'000 |
| Fig. 5b,c; S5a,c | Immunostaining | FLAG | Mouse | Sigma (F1804) | 1:500 |
| Fig. 5c; Fig. S5c | Immunostaining | MuSK | Rabbit | Self made (194T) | 1:8'000 |
| Fig. 5g, Fig. S5a,f | RNAscope + Immunostaining | Laminin | Rabbit | Sigma (L9393) | 1:500 |
| Fig. S5b | Immunostaining | Mouse IgG | Goat | Jackson | 1:500 |
| Fig. S5b | Western Blot | FLAG | Mouse | Sigma (F1804) | 1:1'000 |
| Fig. S5b | Western Blot | alpha-Tubulin | Rabbit | Abcam (ab15246) | 1:800 |
| Fig. S5b | Western Blot | Na/K ATPase | Rabbit | Cell signaling (3010) | 1:1'000 |
| Fig. 6e,f,g | Immunostaining | Synaptophysin (SYP) | Rabbit | Invitrogen (MA5-14532) | 1:300 |
| Fig. 6e,f,g | Immunostaining | Neurofilament (NF) | Rabbit | Sigma (N4142) | 1:15'000 |
| Fig. 7a | Immunostaining | alpha-tubulin | Rabbit | Abcam (ab15246) | 1:200 |
| Fig. 7a,b; S7a,c | Immunostaining | FLAG | Mouse | Sigma (F1804) | 1:500 |
| Fig. 7b; S7c | Immunostaining | GM130 | Rabbit | Cell signaling (D6B1) | 1:3'200 |
| Fig. S7a | Immunostaining | MuSK | Rabbit | Self made (194T) | 1:8'000 |
| Fig. 7c | Western blot | GM130 | Rabbit | Cell signaling (D6B1) | 1:1'000 |
| Fig. 7c,e; S7e | Western blot | MuSK | Rabbit | Self made (194T) | 1:40'000 |
| Fig. 7c,e; S7e | Western blot | FLAG | Mouse | Sigma (F1804) | 1:1'000 |
| Fig. 7e; S7e | Western blot | GAPDH | Rabbit | Cell signaling (14C10) | 1:2'000 |

